# A Large Language Model Guides the Affinity Maturation of Antibodies Generated by Combinatorial Optimization Algorithms

**DOI:** 10.1101/2024.12.19.629473

**Authors:** Faisal Bin Ashraf, Karen Paco, Zihao Zhang, Christian J. Dávila Ojeda, Mariana P. Mendivil, Jordan A. Lay, Tristan Y. Yang, Fernando L. Barroso da Silva, Matthew H. Sazinsky, Animesh Ray, Stefano Lonardi

## Abstract

The ability of an antibody to bind an antigen with high specificity and strength (i.e., its *binding affinity*) are critical properties in the design of neutralizing antibodies. Recent technical advances in AI and a surge of experimental data on antigen-antibody interaction are driving innovations in the design and optimization of antibodies via affinity maturation. Here we introduce Ab-Affinity, a novel large language model which can accurately predict the binding affinity of specific antibodies against a target peptide within the SARS-CoV-2 spike protein. When used in conjunction with a genetic algorithm and simulated annealing, Ab-Affinity can generate novel antibodies with more than a 160-fold increase in predicted binding affinity compared to those obtained experimentally. Our experimental results show that the synthetic antibodies produced by Ab-Affinity have strong predicted biophysical properties. Molecular docking and molecular dynamics simulation of binding interactions of the best synthetic antibodies show enhanced interactions and stability on the target peptide epitope. In general, antibodies generated by Ab-Affinity are superior to those obtained with other existing computational methods.

## 1 Introduction

The sequence motifs that allow an antibody to recognize and bind to an antigen target sequence or *epitope* are embedded within the *complementarity determining regions* (CDR) of the light and heavy chains [1]. The binding of an antigen by the antibody can trigger various immune responses, including the neutralization of a live virus, a toxin, or an antigen present on a diseased cell surface (*e*.*g*., a cancer antigen), leading to the eventual elimination of the antigen or cell from the organism [1]. Direct measurement of the binding affinity of an antibody to its target antigen, which is critical to therapy development, incurs significant manual labor, time, and cost. Computational approaches that use structural databases of known antigen-antibody complexes and machine learning algorithms are increasingly replacing wet lab experiments to narrow the list of candidate antibodies for final laboratory testing [2].

Determining the specificity and strength of an antigen-antibody interaction is a particular case of the problem of predicting whether two proteins can bind to each other [2]. However, the special problem of antigen-antibody binding is more difficult because the epitope and the cognate sequence motif on the antigen (i.e., the *paratope*) are often structurally flexible and contain relatively mobile *intrinsically disordered protein regions*, at least when not bound to each other [3, 4]. Structures of flexible and mobile areas of proteins are poorly represented in protein structure databases because they are technically difficult to determine. In consequence, training data for learning structural rules are sparse. Predicting these interactions from amino acid sequences alone can be advantageous if the structural and functional roles of amino acids in the sequence are effectively identified and modeled.

Predicting accurate epitope-paratope interaction depends in turn on predicting accurate structures of the interacting proteins, which is another computationally hard problem. A major advance occurred when AlphaFold was implemented on the DeepMind architecture [5], using features extracted from amino acid sequences of proteins combined with known structural information, and from multiple sequence alignments of evolutionarily related protein sequences. AlphaFold2 achieved even more remarkable accuracy [6], comparable to experimentally determined structures, and garnered widespread attention for its transformative impact on structural biology. Capitalizing on the successful prediction of single protein structures, Alphafold-Multimer could predict structures of interacting proteins with modest success [7]. Shortly thereafter, Alphafold3 vastly improved prediction by incorporating a diffusion module for approximating atomic positions, besides re-framing the representational learning of physicochemical properties of proteins and their evolutionary relationships from multiple sequence alignments [8]. While Alphafold3 significantly improved the prediction accuracy of multi-chain protein complexes, the accuracy of predicting antigen-antibody complexes has been limited, producing about 60% correct prediction and 30-40% high accuracy prediction for antibodies with low sequence homology to known structures. These successes soon led to attempts to learn by natural language processing the “grammatical rules” by which polypeptide chains spontaneously fold on natural protein sequence strings, which evidently have been selected by evolution for optimized function. In this framework, the amino acids form the alphabet, short amino acid stretches are words, and entire sequences are sentences in which the probability distribution of the immediate neighbors of an amino acid within the polypeptide string is guided by probabilistic rules that could have been “shaped by evolution”. Stochastic context-free grammars were also used to capture long-distance dependencies in amino acid positions [9–12]. The first successful Large Language Model (LLM) for predicting protein structure, ESMfold, used an unsupervised transformer model with about 15 billion parameters, which was trained on the sequence of 138 million proteins [13]. More recent models such as DG-Affinity [14] and CSM-AB [15] integrate LLMs and graph-based techniques, thus enhancing the accuracy of the prediction of antigen-antibody interaction. Tag-LLM [16] and FAbCon [17] further refined these capabilities using task-specific tags and fine-tuned LLMs. The use of pre-trained protein language models in DG-Affinity [14] has shown superior performance in predicting antibody binding affinities compared to several other methods. LLMs were recently used to predict the binding affinity of natural antibodies against the SARS-CoV-2 spike protein [18].

Recent advances in generative AI have made significant progress in the *de novo* design of proteins (including antibodies) that perform new functions (see *e*.*g*. [19, 20]). Protein structure-based diffusion models have been used to optimize sequence and structure simultaneously, to achieve the target function [21]. Several LLMs have shown remarkable performance in capturing protein properties directly from sequences [22]. For example, ProtGPT2 [11] and ProGEN2 [12] are auto-regressive language models trained on protein sequences that can generate new proteins with a desired function. ESMDesign [23] employs a masked large language model combined with simulated annealing optimization and ESMFold-based scoring to refine protein designs. ESMFold includes the large language model ESM-2, which was trained on the UniRef50 database [13]. Diffusion models such as RFDiffusion [24] and EvoDiff [25] offer more refined controls. RFDiffusion allows conditioning on attributes such as chain length and binding characteristics, while EvoDiff focuses on multiple sequence alignment families for evolutionary consistency. Generative AI is also revolutionizing the process of designing therapeutic antibodies (see e.g. [26–30]). Hybrid approaches that employ both laboratory methods and ML algorithms have been tested (see e.g. [2, 31–39]). RefineGNN [40] uses graph neural networks trained on antibody sequences and structures, while IgLM [41] and DiffAb [21] provide enhanced control through conditioning on chain type, species, and specific target antigens. In a recent study, the binding affinity landscape of a library of mutant antibody sequences was generated using an LLM, demonstrating optimized binding properties of the synthetic antibodies [18]. Notably, general Protein Language Models can guide affinity maturation of antibodies using evolutionary information alone and can achieve competitive performance compared to *in vitro* methods [42].

Here, we demonstrate for the first time, to the best of our knowledge, the generation of novel antibody molecules with improved binding characteristics against a target antigen using a genetic algorithm and simulated annealing, both guided by an LLM specifically trained on antigen-antibody binding data.

## 2 Results

### 2.1 Overview of the computational framework

Our strategy began with a population of natural antibody sequences, including the heavy and light chains and linker sequences (as illustrated in Figure 1). These sequences were used to train Ab-Affinity to predict embeddings, binding affinity, and intra-residue attention maps. To optimize the antibody design, we employed genetic algorithms and simulated annealing. The genetic algorithm starts with an initial population of antibodies, then iteratively applies selection, crossover, and mutation to identify the best-performing candidates. Simulated annealing begins with a seed antibody, then generates and evaluates its nearest neighbors iteratively, and selects candidates that improve affinity by gradient descent. These processes ultimately converged on locally optimal candidate antibodies against the HR2 peptide of the SARS-CoV-2 spike protein.

**Figure 1:**
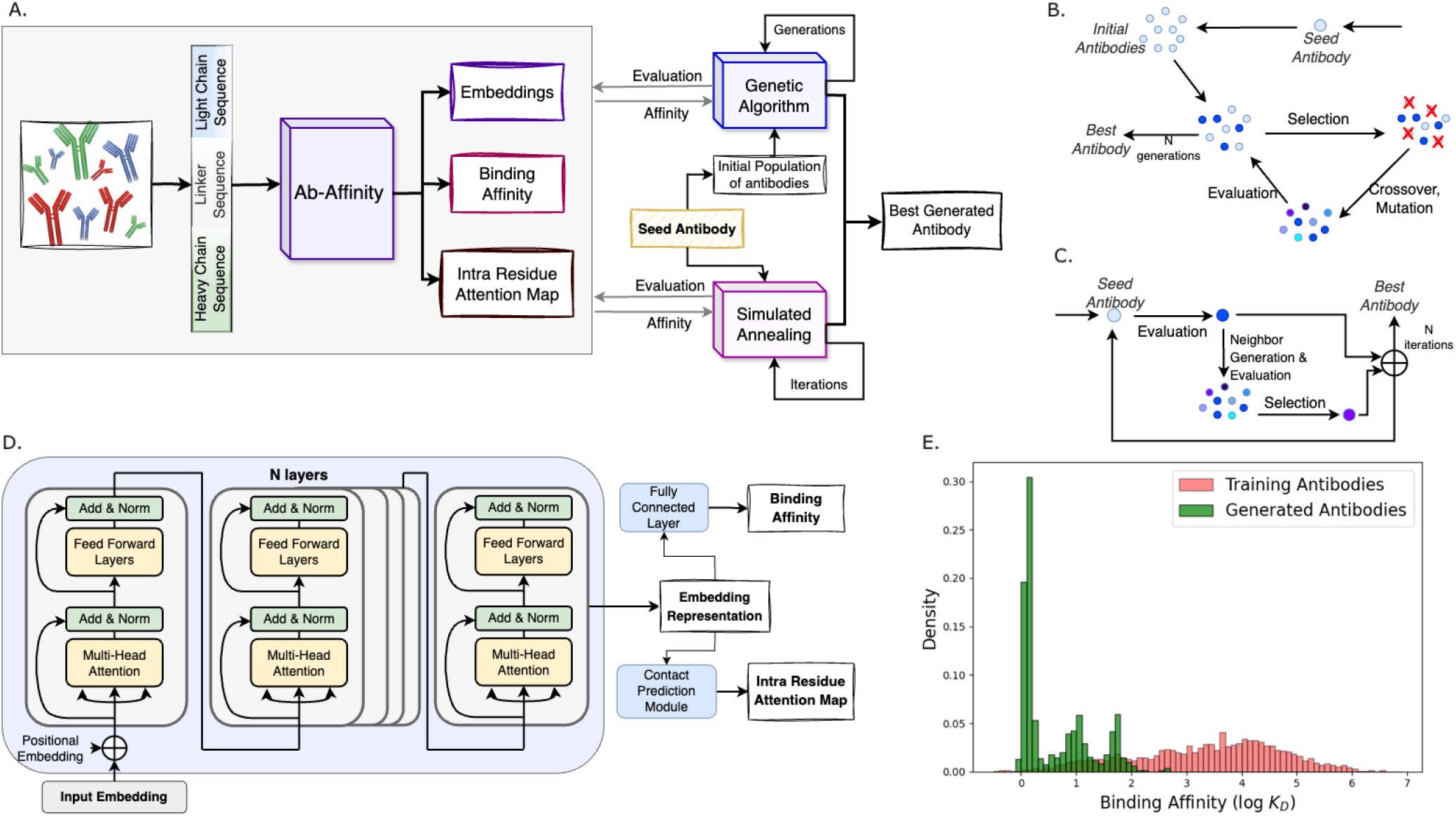
Overview of the framework for generating and optimizing antibody binding affinity. **A**. Ab-Affinity predicts the binding affinity against a specific target peptide; it also provides residue-residue contact maps and an embedding of the input sequence; a genetic algorithm and a simulated annealing method, in combination with Ab-Affinity, generate new synthetic antibodies. **B**. The genetic algorithm generates synthetic antibodies starting from a seed antibody by introducing mutations and crossover operations, guided by the predicted affinity scores from Ab-Affinity. **C**. The simulated annealing method starts from a seed antibody and explores its sequence neighbors, guided by the predicted affinity scores from Ab-Affinity. **D**. The architecture of Ab-Affinity has *N* sequential layers, each containing multi-head attention, which constitute the encoder of the model that generates the embedding representation; the fully connected layer and the contact prediction module are on top of the encoder to predict binding affinity and residue-residue attention map. **E**. Binding Affinity distribution of synthetic antibodies and training antibodies; synthetic antibodies (green) have stronger binding affinities (lower log *K*_*D*_) than training antibodies (red); all *K*_*D*_ values considered in this work are in units of 10^−9^*M* (*nM*).

### 2.2 Ab-Affinity Predicts the Binding Affinity of Antibodies Against the HR2 Domain of SARS-CoV-2 Spike Protein

First, we investigated whether Ab-Affinity generates meaningful latent space features (*i*.*e*., embeddings) for the antibodies. We compared Ab-Affinity’s ability to predict binding affinity with those of three other LLM-based methods, namely DG-Affinity, ESM-2, and AbLang. The same dataset, which was held out from the training and was not utilized during model development, was used to compare the predicted binding affinities against experimentally observed values by all three methods. We should note that (i) DG-Affinity uses a pre-trained language model combined with a ConvNext-based architecture for prediction, which outperformed 26 other methods on an independent antibody dataset [14], (ii) ESM-2 was pre-trained on the UniRef50 protein database and generated 640-dimensional antibody sequence embeddings, which were fed into a simple linear regression model, and (iii) AbLang uses separate embeddings for the heavy and light chains of antibodies, which were concatenated into a 1534-dimensional vector and used in a linear regression model to predict the affinity score.

The results are summarized in Figure 2. Observe that Ab-Affinity’s predictions produced the highest Pearson and Spearman correlation coefficients. The scatter plot in Figure 2B shows the distribution of predicted log *K*_*D*_ values relative to the experimentally determined values. DG-Affinity exhibited the lowest Pearson correlation coefficient, presumably because despite using an LLM pre-trained on antibodies, the architecture was not trained with any SARS-CoV-2-specific data. AbLang, trained on antibody sequences, also exhibited high Pearson/Spearman correlation coefficients. ESM-2, pre-trained on ‘generic’ proteins, showed slightly higher Pearson/Spearman correlations than AbLang but performed worse than Ab-Affinity. The scatter plots in **Supplementary Figure 4** of actual *vs*. predicted binding affinity show that the best-fit lines for Ab-Affinity have the highest slope, indicating the best predictive accuracy. The Pearson correlation coefficient measures the linearity of the relationship between the observed and predicted values, which is sensitive to outliers. Pearson’s rank correlation is insensitive to outliers and is robust to nonlinear and monotonic trends. The scatter plots show a relative absence of extreme outliers in the test dataset, suggesting the linearity of prediction with observed values by all three methods.

**Figure 2:**
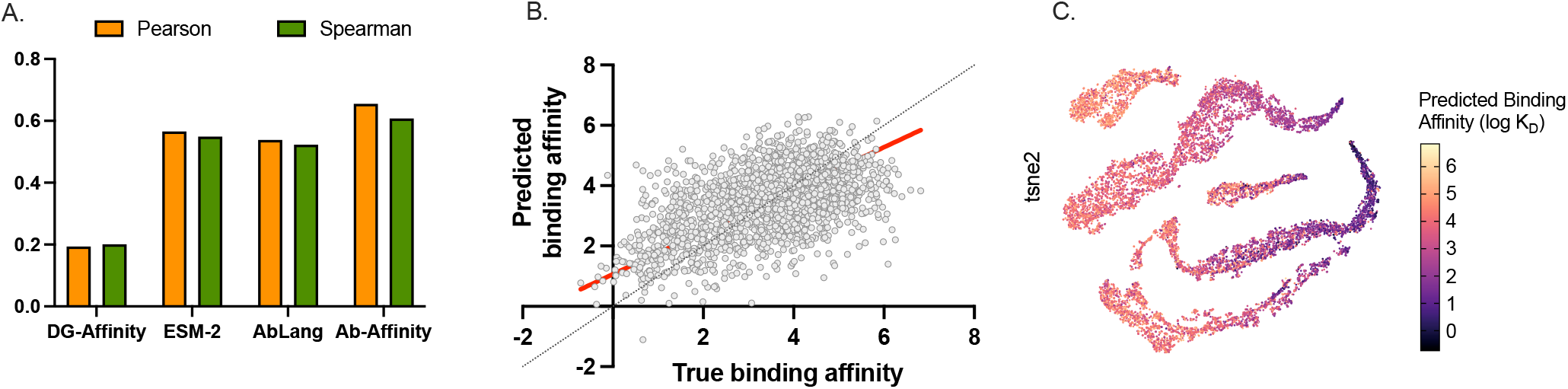
Performance of Ab-Affinity, ESM-2, AbLang and DG-Affinity on predicting the binding affinity of antibodies against the HR2 domain of SARS-CoV-2 spike protein. **A**. Pearson and Spearman correlation for DG-Affinity (p-values = 3.86 × 10^*−*14^, and 3.38 × 10^*−*15^), ESM-2 (p-values = 8.03 × 10^*−*198^, and 8.02 × 10^*−*198^), AbLang (p-values = 1.24 × 10^*−*175^, and 1.02 × 10^*−*163^) and Ab-Affinity (p-values = 4.03 × 10^*−*261^, and 8.65 × 10^*−*217^). **B**. Scatter plot of actual *vs*. predicted binding affinity using Ab-Affinity. **C**. t-SNE representation of the embedding produced by Ab-Affinity; antibodies are colored according to their binding affinity log *K*_*D*_ (*K*_*D*_ was in units of 10^*−*9^ M).

We visualized Ab-Affinity’s embeddings in 2D space using t-distributed Stochastic Neighbor Embedding (t-SNE) maps. To evaluate Ab-Affinity’s performance, we also visualized the embeddings produced by ESM-2. In the t-SNE plots in Figure 2C each point is an antibody and the color illustrates its corresponding binding affinity. As shown in Figure 2C, the embeddings produced by Ab-Affinity display a smooth gradient of binding affinities, *i*.*e*., the value of log *K*_*D*_ monotonically decreases. By contrast, the ESM-2 embeddings in **Supplementary Figure 3** do not show a clear gradient along the t-SNE components, suggesting that the embeddings produced by Ab-Affinity are more informative to capture antigen-antibody interactions.

To explore whether the prediction methods are sensitive to the choice of the test dataset, we compared the performance of Ab-Affinity with those of the other three methods using a different dataset. Table 1 compares the performance of CNN-based and LLM-based methods for predicting binding affinity to the same peptide. We used the 14H dataset (heavy chain sequences) and 14L dataset (light chain sequences), which are the mutated versions of Ab-14 heavy and light chain sequences, respectively. To fit the heavy chain sequences from the 14H dataset, the seed light chain sequence of Ab-14 was paired with different heavy chains from 14H to form scFv sequences. Similarly, the heavy chain seed was paired with different light chains from 14L to form additional scFv sequences. Table 1 shows that the Pearson correlation coefficient of both 14H and 14L antibodies for Ab-Affinity was the highest among all models. While the highest Spearman correlation coefficient for 14H was obtained by A2Binder, that of Ab-Affinity closely followed. Scatter plots for actual *vs*. predicted binding affinity for these methods on both 14H and 14L datasets are given in **Supplementary Figures 5 and 6**. Note that these correlation coefficients are specific to predicting the binding affinity for antibodies derived from the Ab-14 seed antibody. However, predictions from Ab-Affinity not only showed a higher correlation coefficient for Ab-14 but also for all three seed antibodies.

**Table 1:** Comparing Ab-Affinity against other binding affinity predictors on the 14H and 14L data sets; numbers in bold-face indicate the best performance; N/A indicates that the tool is not applicable for the dataset or that performance measure is not available from the corresponding paper

| Model | Ref | 14H |  | 14L |  |
| --- | --- | --- | --- | --- | --- |
|  |  | Pearson | Spearman | Pearson | Spearman |
| Ens-Grad | [33] | 0.601 | 0.476 | 0.637 | 0.645 |
| ESM-F | [43] | 0.634 | 0.516 | 0.674 | 0.681 |
| AntiBERTa2 | [44] | 0.623 | 0.545 | 0.673 | 0.684 |
| AbMAP | [45] | 0.606 | 0.510 | 0.674 | 0.685 |
| A2Binder | [43] | 0.642 | **0.553** | 0.683 | 0.688 |
| Ensembles-14H | [18] | N/A | 0.512 | N/A | N/A |
| Ensembles-14L | [18] | N/A | N/A | N/A | 0.688 |
| **Ab-Affinity** | [this] | **0.652** | 0.526 | **0.712** | **0.713** |

The closest performance on this training set came from A2binder [43], a model that predicts antibody binding affinity against various SARS-CoV-2 spike protein epitopes. However, there are key differences between A2binder and Ab-Affinity. Methodologically, A2binder uses three separate pre-trained language models for the antibody heavy chain, light chain, and antigen epitope, followed by a complex CNN module. In contrast, Ab-Affinity relies on a single language model without additional modules. Additionally, A2binder requires the antigen sequence for each SARS-CoV-2 variant, whereas Ab-Affinity targets a conserved peptide present in all variants. Experimentally, Ab-Affinity offers richer interpretability and utility. It generates meaningful antibody embeddings that capture both binding properties and structural robustness, enabling visualization of the antibody binding landscape. Furthermore, it provides intra-residue attention maps, offering insights into residue-residue interactions that distinguish strong from weak binders.

### 2.3 Ab-Affinity’s Embeddings Enable Downstream Classification Tasks

We used the embeddings produced by Ab-Affinity to carry out two downstream classification tasks, namely (i) the problem of determining the antibody binding affinity classes (High, Medium, Low), and (ii) the problem of determining whether the binding affinity of an antibody was stronger than the binding affinity of the corresponding seed antibody.

In the first task, Ab-Affinity achieved an AUC of 0.92 for strong antibody classification, significantly outperforming ESM-2’s AUC of 0.78. For weak antibodies, the AUCs were 0.78 and 0.71 for Ab-Affinity and ESM2, respectively (see also **Supplementary Figure 7**). In the second task, Ab-Affinity demonstrated superior performance with an AUC of 0.91 relative to ESM-2’s 0.74 (see also **Supplementary Figure 7**). These results suggest that Ab-Affinity’s embeddings are more informative for antigen-antibody binding optimization.

### 2.4 Ab-Affinity’s Attention Maps for Strong and Weak Binding Antibodies

We used attention maps obtained from Ab-Affinity to determine which residue-residue interactions were more important for the prediction of binding affinity. We extracted and compared two residue-residue contact maps, namely (i) heavy and light chains with strong binding affinity (i.e., log *K*_*D*_ *<* 0.5) and (ii) heavy and light chains with weak binding affinity (i.e., log *K*_*D*_ *>* 5.5). **Supplementary Figure 8** illustrates the residue-residue differences between the attention maps for strong and weak binding antibodies for both heavy and light chains, where the CDRs are highlighted. Observe that many of the strongest differences occur in CDR-H1, CDR-H2, CDR-L1, or the regions immediately adjacent to them. Figure 3A shows the differences in Ab-Affinity’s attention between a weak- and a strong-binding antibody. Observe that the significant difference in attention in the CDR regions. Figures 3B and 3C map the residue-residue attention relationships in their corresponding 3D structure derived using ESMFold. Observe that residues M34, Y167 and Y170 in the CDR are attending to more residues in the stronger antibody than in the weaker antibody. For the stronger antibody, most of the attended residues are distant from the CDR residues, reflecting a “geometrical need” for binding to the target epitope at a distance.

**Figure 3:**
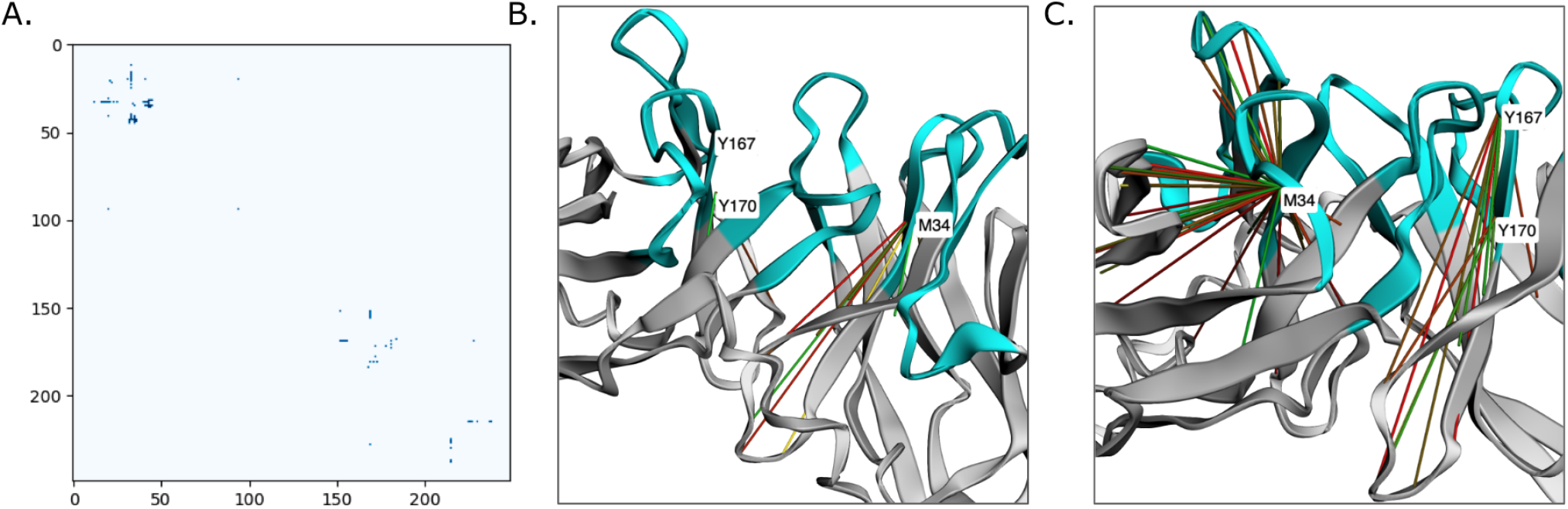
**A**. Differences in residue-residue attention matrix from Ab-Affinity between a weak binding antibody AAYL51 11847 with log *K*_*D*_ = 6.816 and a strong binding antibody Ab-91-SA-pssm-1 with log *K*_*D*_ = 0.799 for their scFv sequence input. Both antibodies were derived from seed Ab-91; **B**. Residue-residue pairs (solid lines) with high attention values mapped in 3D structure of the weak antibody; **C**. Residue-residue pairs (solid lines) with high attention values mapped in 3D structure of the strong antibody; the color of the solid lines indicates the corresponding residue-residue distances in the structure, with red being a long distance (residues are far away) and green being a shorter distance (residues are in close contact); light blue regions are the CDRs. All *K*_*D*_ were in units of 10^−9^ M).

### 2.5 The Thermal Stability Feature Co-Clusters with Ab-Affinity Embeddings

Robustness to changes in temperature is important for therapeutic antibodies. Whereas thermal stability (thermostability) was not an explicit feature on which Ab-affinity was trained, we investigated whether the embeddings produced by Ab-Affinity exhibited a correlation with thermostability. We used as a reference a dataset of experimentally determined thermostability of 26 antibodies against SARS-CoV-2 [42, 46]. In the 2D visualization of the antibody embeddings in **Supplementary Figure 9**, each point is an antibody, and the color illustrates its corresponding experimental thermostability value. In Ab-Affinity’s embedding, the antibodies are separated into two distinct clusters, each with relatively similar thermostability values. This is not the case for the ESM-2 embeddings, where the clusters display mixed values of thermostability.

### 2.6 Ab-Affinity’s Synthetic Antibodies have Stronger Binding Affinity

We explored eight approaches to generate synthetic single-chain variable fragment sequences. The Uniform method generated random amino acids selected with uniform probability at random positions, also selected with uniform probability. In the PSSM method, we used a position-specific scoring matrix to drive the selection of amino acids. The other six approaches fall into two groups. In the first group, we employed a genetic algorithm that can either use (i) random mutations with uniform probability, (ii) random mutations based on the probability distribution in the PSSM, and (iii) random mutations based on the ABModel matrix [47]. In the second group, we used a simulated annealing strategy, again using all three mutation techniques (i-iii) above. Table 2 reports the main statistics for the best-performing synthetic antibodies generated by the eight methods. For each method, we began with one of the three seed antibodies: Ab-14, Ab-91, or Ab-95. The table shows the predicted binding log *K*_*D*_ and the fold improvement relative to the seed antibody. Not surprisingly, the Uniform method fails to produce any meaningful improvement in binding affinity and serves as the baseline. The PSSM method showed substantial improvements in binding affinity. For instance, Ab-14’s binding improved by approximately 13-fold, Ab-91 by 3.5-fold, and Ab-95 by 24-fold. The methods based on the Genetic Algorithm or Simulated Annealing yielded more substantial improvements. The GA-ABModel method for Ab-95 reached a 147-fold improvement, corresponding to ΔΔG = –2.95 kcal/mol relative to the seed antibody, and the SA-Random method achieved the highest (168-fold) enhancement, corresponding to ΔΔG = –3.3 kcal/mol or 1-3 additional hydrogen bonds.

**Table 2:** Predicted binding affinity and improvement over the seed antibody of the top three synthetic antibodies generated by various methods within Ab-Affinity (bold font indicates the best results; all *K*_*D*_ were in units of 10^*−*9^ M); fold improvements were rounded to the nearest integer.)

|  | Ab-14 |  | Ab-91 |  | Ab-95 |  |
| --- | --- | --- | --- | --- | --- | --- |
| | Predicted Binding<br>(log $K_D$) | Fold Improvement | Predicted Binding<br>(log $K_D$) | Fold Improvement | Predicted Binding<br>(log $K_D$) | Fold Improvement |
| Seed Antibody | 1.39 | 1 | 2.45 | 1 | 2.22 | 1 |
| Uniform | 4.25 | - | 3.36 | - | 3.29 | - |
| PSSM | 0.28 | 13 | 1.91 | 3 | 0.85 | 24 |
| GA - Random | 0.14 | 18 | 0.86 | 39 | 0.04 | 150 |
| GA - PSSM | 0.13 | 19 | 1.75 | 5 | 0.72 | 31 |
| GA - ABModel | 0.12 | 19 | 0.98 | 29 | 0.05 | 147 |
| SA - Random | 0.13 | **19** | 0.85 | 40 | -0.01 | **168** |
| SA - PSSM | 0.16 | 17 | 0.79 | **45** | 0.04 | 150 |
| SA - ABModel | 0.13 | **19** | 1.39 | 12 | 0.03 | 154 |

Figure 4A shows density plots of predicted binding affinity (log *K*_*D*_) for synthetic antibodies generated from seed antibody Ab-91 using GA and SA methods. For comparison, we also show the binding affinity distributions for antibodies from the training set, PSSM-based generation, and Random substitution. As expected, Ab-Affinity improved the affinity of seed antibodies. In contrast, the PSSM and Random methods showed little to no improvement over the training set (with similar median values). Additionally, for all seed antibodies (Ab-14, Ab-91, Ab-95), the density plots indicated that all generation methods produce antibodies with improved predicted affinity compared to the training libraries, as reflected by lower median log *K*_*D*_ values (see also **Supplementary Figure 10**).

**Figure 4:**
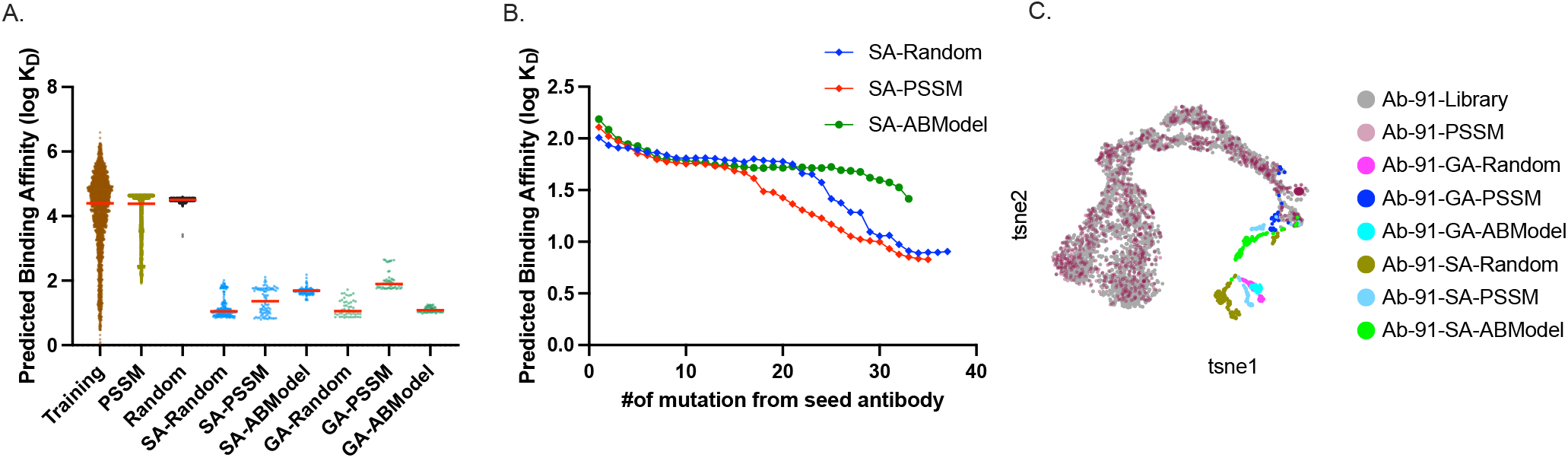
Analysis of generated antibodies from seed antibody Ab-91. **A**. Distribution of predicted binding affinity for the antibodies in the training library compared to the synthetic antibodies. The red horizontal line indicates the median of the distribution; **B**. Predicted binding affinity of the synthetic antibodies as a function of the number of mutations from the corresponding seed antibody; **C**. t-SNE projection for the Ab-affinity embedding of the antibodies in the training library and the synthetic antibodies obtained from the Ab-91 seed. All *K*_*D*_ were in units of 10^*−*9^ M).

Next, we investigated how the different approaches explored the sequence landscape within the same number of trials. To this end, we computed the mutational distance of synthetic antibodies that had a stronger binding affinity than the seed sequence. We found that some generative methods were more conservative than others in exploring the fitness landscape. For example, the PSSM approach generated antibodies with up to 9 mutations, whereas the genetic algorithm had a broader exploratory reach, spanning mutational distances of 24, 34, and 37 for Ab-14, Ab-91, and Ab-95, respectively. Simulated annealing generated even more diversity, producing sequences up to 26, 37, and 44 mutational distances for the respective antibodies (see **Supplementary Figure 11**). We analyzed the relationship between the number of mutations and the predicted binding affinity using simulated annealing to estimate how the exploratory space affects selection. For Ab-14 and Ab-95, the binding affinity improved significantly as more mutations were explored. In the case of Ab-91 (see Figure 4B), only SA-Random and SA-PSSM showed consistent improvements in binding affinity as the number of mutations increased. Interestingly, binding affinity remained relatively stable until a threshold number of mutations was reached, after which it improved markedly, suggesting cooperative events. In the case of SA-AbModel, this initial plateau phase was more prolonged.

### 2.7 Ab-Affinity’s Synthetic Antibodies are Novel

To compare the synthetic antibodies to antibodies in the original library, we projected the embeddings in a 2D space using t-SNE (Figure 4C and **Supplementary Figure 12**). For example, in the t-SNE plot for Ab-91 in Figure 4C the synthetic antibodies generated by GA-Random, GA-PSSM, GA-ABModel, SA-Random, SA-PSSM, and SA-ABModel formed distinct clusters that were clearly separated from the library antibodies (yellow dots). Similar observations can be made for Ab-14 and Ab-95 in **Supplementary Figure 12**. We should note that our approaches generate synthetic antibodies with up to 44 mutations from the seed sequence, whereas the training dataset contains antibodies with up to three mutations. Our methods can explore much larger subspaces of the sequence diversity landscape, generating candidates whose sequence is far from the antibodies in the original library. As expected, the optimized synthetic antibodies produced by our methods have binding affinities significantly higher than those in the training library (see **Supplementary Figure 12**).

### 2.8 Ab-Affinity’s Synthetic Antibodies have Optimized Biophysical Properties

Next, we analyzed the biophysical properties of synthetic antibodies, which are crucial to their overall performance against a target antigen. The isoelectric point (pI) influences antibody-antigen interaction by determining the pH at which these macromolecules carry a net charge of zero. At pH values different from the pI, the net charge of the antibody affects its binding kinetics with the antigen. Human antibodies typically have pI 7.5-9 [48]. Synthetic antibodies generated using GA-Random, GA-ABModel, SA-ABModel, and SA-PSSM from the Ab-14 seed also had pI 7.5-9.0 (**Supplementary Figure 13**). From the Ab-91 seed, all GA and SA antibodies had a pI higher than 7.0. The hydrophobicity index determines solubility; synthetic antibodies generated by SA were less hydrophobic compared to library antibodies for all three seed antibodies, suggesting that the former antibodies are likely to be more soluble. The pattern of *hydrophobicity* of a protein, especially on its surface, influences its thermal stability and solubility - the lower the surface hydrophobicity, the more stable and soluble [27]. The *instability index* measures the overall stability of the protein structure. An instability index higher than 40 indicates instability [49]. Our synthetic antibodies exhibited a lower instability index (lower than 40) compared to library antibodies. Finally, the secondary structure provides insights into the antibody’s binding abilities. For example, efficient binding is expected for antibody variable regions that contain a higher content of polypeptide backbone turns, rather than possessing mostly alpha helices and/or beta sheets, suggesting the presence of flexible and globular regions. The higher backbone turn contents of our synthetic antibodies compared to library antibodies, relative to the proportion of *α*-helices and *β*-sheets, suggest better structural properties of their interaction surfaces relative to those of the starting library antibodies (**Supplementary Figure 14**).

### 2.9 Structural Docking and Molecular Dynamics Simulation Validates the Top Synthetic Antibodies

We first carried out molecular docking to assess the interaction of our synthetic antibodies with the target antigen. We used ESMFold to obtain the predicted 3D structures of our synthetic antibodies, which were docked to the SARS-CoV-2 spike protein using HADDOCK. HADDOCK generated several docking poses corresponding to different feasible conformations, which were clustered based on energy and physical properties. For each antibody—comprising one seed and candidates from six different approaches—multiple antigen-bound conformations were produced. These were clustered, and the cluster with the best HADDOCK score (a weighted combination of various energy terms) was selected to represent each antibody–antigen complex. This yielded a single representative docking score per antibody. These scores, along with their associated variations, were then used to calculate p-values, which are summarized in Figure 5. A lower HADDOCK score indicates higher complex stability. With one exception (SA for Ab-95), all synthetic antibodies were predicted to form more stable complexes than the seed antibody (Figure 5A). We computed the *buried surface area* inaccessible to the solvent, due to complex formation (which is directly proportional to 1*/K*_*D*_, [50]) of each antigen-antibody complex. As anticipated, all synthetic antibodies except one had higher buried surface areas than the corresponding seed antibody when bound to the antigen (Figure 5B). This increase in buried surface area suggests more specific binding interactions, as greater contact between proteins generally correlates with enhanced specificity and affinity in protein-protein interactions [51, 52]. Note that HADDOCK scores for different docked complexes are not directly comparable; however, consistently better HADDOCK scores of the generated antibodies against the target HR2 epitope support the argument that the optimized antibodies might indeed have better binding characteristics.

**Figure 5:**
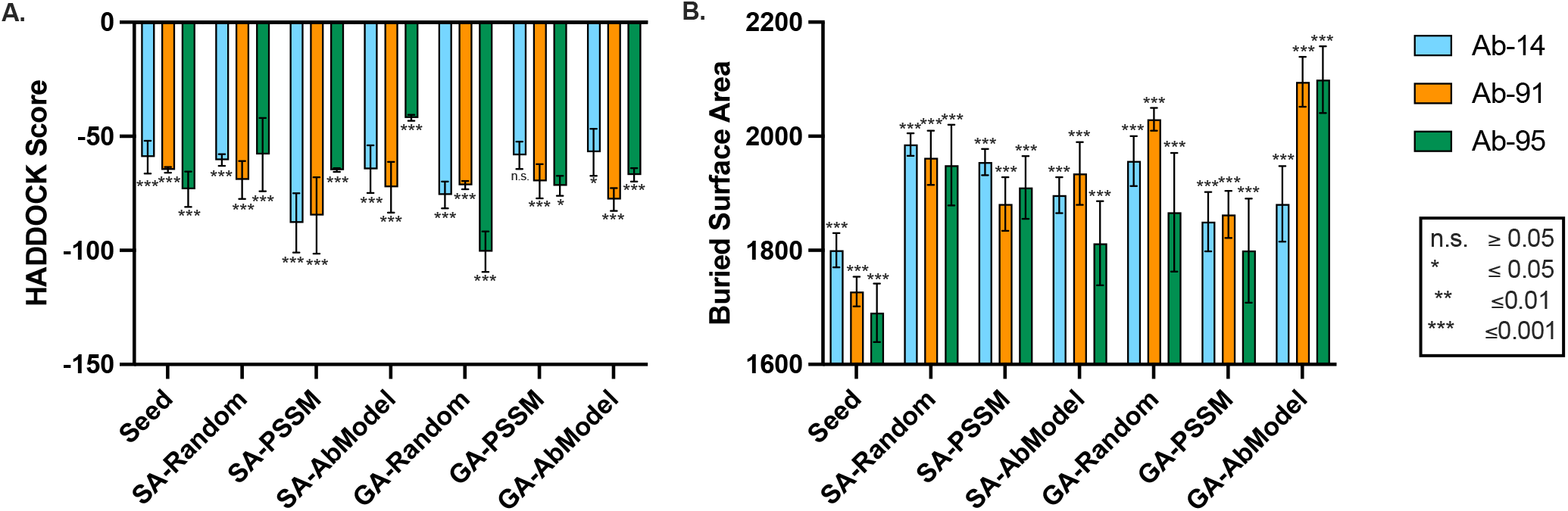
Molecular Docking of the antibodies with SARS-CoV-2 spike protein. **A**. Average HADDOCK-Score for the best synthetic and seed antibody [lower score indicates stronger binding]; **B**. Average buried surface area [higher the better] for the best synthetic and seed antibody; the error bar indicates the standard deviation; significance p-values compared to seed antibody are shown by the asterisks;

To confirm optimized complex formation, we conducted MD simulations up to 100 ns for the complexes containing the predicted Ab-14-SA-PSSM-1, Ab-14-SA-PSSM-6, and their seed Ab-14-seed, and we analyzed the trajectories of CDRs as well as the target epitope residues, respectively, during the final 100 ns of the simulation. In each case, we started with the highest-scoring docked complex; the residue-averaged root mean square deviation (RMSD) values oscillated within the first 40 ns, but stabilized by 100-200 ns (Supplementary Figure 16). Whereas the aggregate RMSD value in the final 100 ns was significantly reduced for the epitope residues, we did not observe a similar trend with the CDR residues. Note that the target epitope was a contiguous stretch of 14 amino acids near the N-terminus of a 55-residue long HR2 region that is mostly alpha-helical. On the other hand, the CDR region is substantially longer (63 amino acids), contains several flexible regions, and only a minority of the CDR residues is expected to be in contact with the epitope. Once the trajectories stabilized, the mobility of the epitope atoms was significantly restrained in complexes with the optimized antibodies relative to those with the corresponding seed antibodies. These observations held also with the Ab-95-seed and its optimized derivatives, suggesting that the optimization algorithm has learned certain general physical rules governing epitope-paratope interaction.

In the complex containing Ab-14-SA-PSSM-1, the HR2 epitope was stabilized by a network of interactions (Figure 6A) involving two charged residues of the epitope (ASP9 and ASP12), and four CDR3 residues (ARG100, SER101, ALA102 and LYS98), together potentially contributing –19.7 to –26.9 kcals/mole of interaction energy (Table 3). Similarly, the complex containing Ab-14-SA-PSSM-6 is stabilized by interactions (Figure 6B) between four epitope residues (GLY11, ASP12, SER14 and ASN17) and two CDR3 heavy chain residues (THR101 and LYS98), potentially contributing –8.5 to –14.0 kcals/mole of interaction energy. Interestingly, in each case of the optimized antibody-epitope complex, there was significantly higher (relative to the unoptimized seed) contribution of electrostatic interaction energy, but also a complementary solvation energy penalty (Figure 7A, D). The higher contribution of charge-charge interaction in the epitope-paratope interactions (Figure 7B-C,E-F) is consistent with previous studies [55]. Furthermore, neutralization of key ionizable residues at the antigen-antibody interface (namely ASP90, ASP106, ARG100, and LYS98 on the antibody side, and ASP7, ASP9, and ASP12 on the antigen) led to a decrease in binding affinity, as evidenced by the constant-pH CG simulations (see Table 2 in the **Supplementary Materials**).

**Table 3:** Predicted residue-residue interactions between optimized antibodies and peptide HR2. The table lists the interacting residue pairs along with their interatomic distances (in A°) and the dominant interaction type, as predicted from molecular dynamics simulations.

| Optimized Antibody | Interacting Amino Acid Residue |  | Distance (Å) | Interaction Type |
| --- | --- | --- | --- | --- |
|  | in HR2 | in antibody |  |  |
| **Ab14-pssm1** | ASP9 | ARG100 | 2.817 | Salt bridge, Hydrogen bond |
|  | ASP12 | ARG100 | 2.768 | Salt bridge, Hydrogen bond |
|  | ASP12 | SER101 | 2.631 | Electrostatic: Ion-dipole, Hydrogen bond |
|  | ASP12 | ALA102 | 4.905 | Induction + dispersion |
|  | ASP9 | LYS98 | 2.792 | Salt bridge, Hydrogen bond |
| **Ab14-pssm6** | GLY11 | THR101 | 3.116 | Salt bridge, Hydrogen bond |
|  | ASP12 | LYS98 | 4.043 | Salt bridge, Hydrogen bond |
|  | ASP12 | THR101 | 2.718 | Electrostatic: Ion-dipole, Hydrogen bond |
|  | SER14 | THR101 | 3.739 | Electrostatic: Ion-dipole, Hydrogen bond |
|  | ASN17 | THR101 | 3.576 | Electrostatic: Ion-dipole, Hydrogen bond |

**Figure 6:**
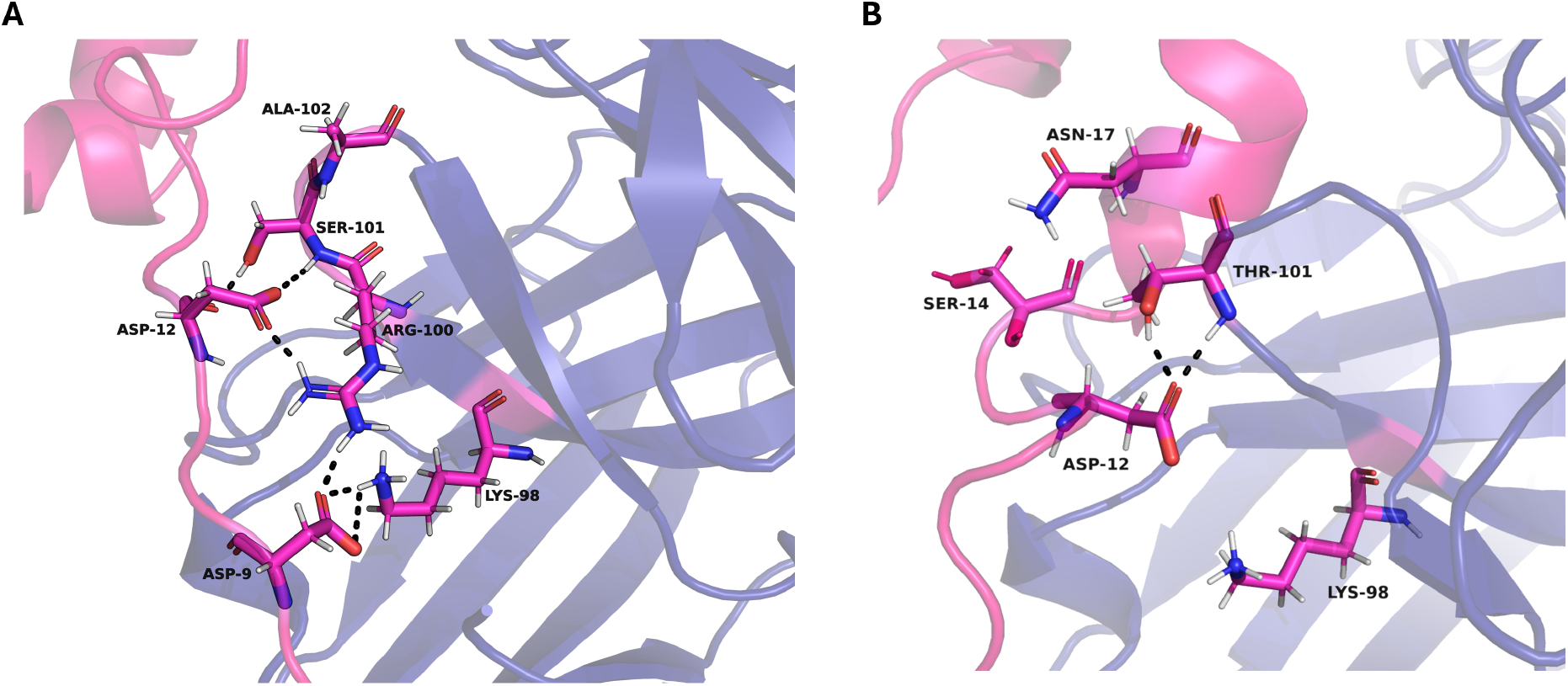
Optimized antibodies make stable complexes. MD simulations were run for 100 ns, and the contact residues at the CDR3 are shown. Complex of HR2 epitope with Ab-14-SA-PSSM-1 (**A**.), and with Ab-14-SA-PSSM-6 (**B**.). Contact residues were calculated with MAPIYA [53] and atomic distances were displayed using PyMOL [54] (PyMOL version 3.1.4).

**Figure 7:**
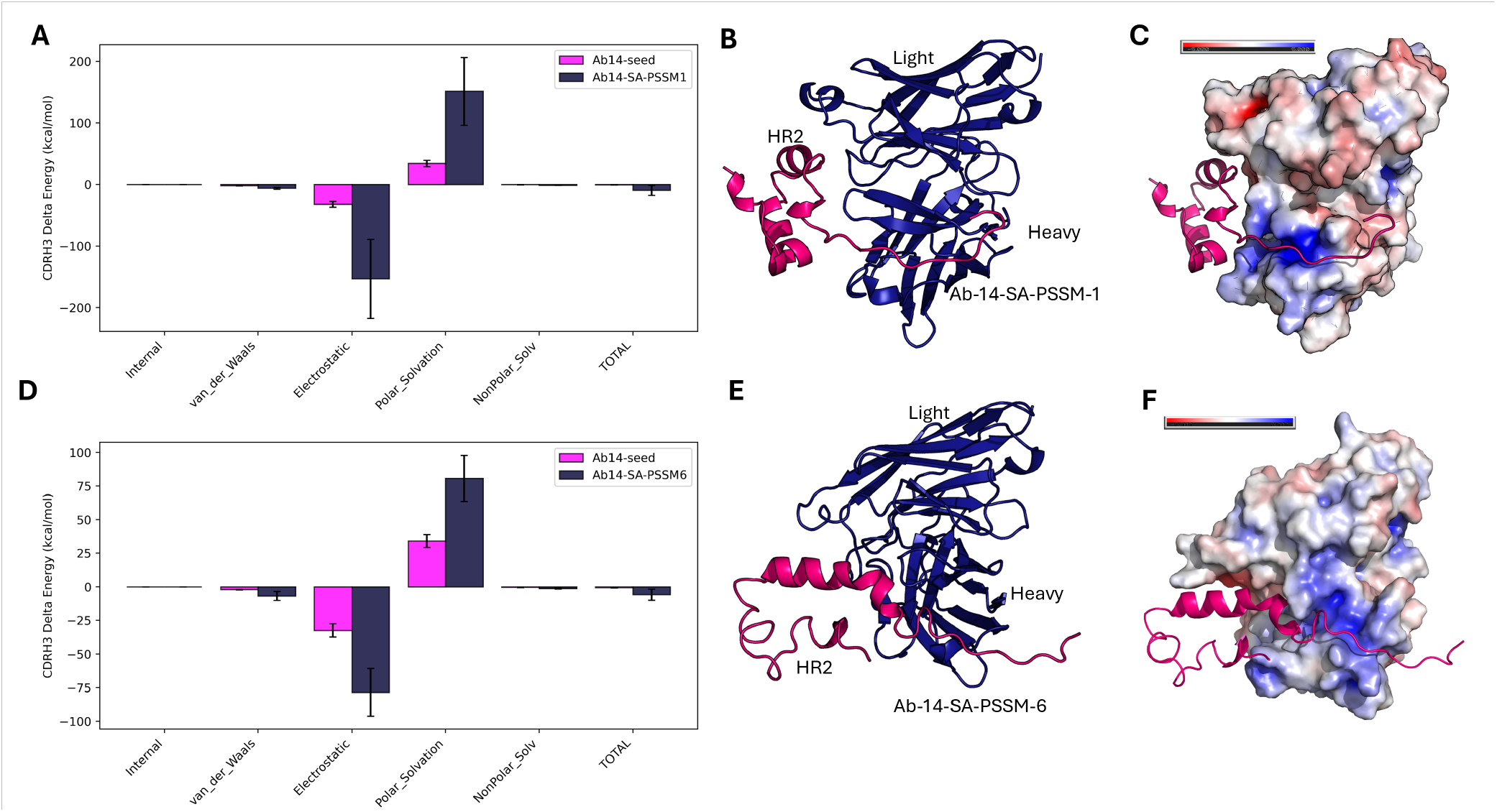
Molecular Dynamics simulation of the antibodies in complex with the HR2 region of SARS-CoV-2 spike protein. **A, D**. Binding free energy from MM-PBSA at the CDR3 heavy chain (red: Optimized antibodies Ab-14-SA-PSSM-1 and Ab-14-SA-PSSM-6; blue: Ab-14 seed). **B, E**. Representative three-dimensional structure of the top cluster from MD trajectories shows preferential binding of HR2 for the heavy chain of the optimized antibodies (pink:HR2, dark-blue: Optimized antibodies Ab-14-SA-PSSM-1 and Ab-14-SA-PSSM-6). **C, F**. The Electrostatic potential of the antibodies using APBS (PyMOL version 3.1.4) mapped at the antibody surfaces indicates the strong Coulombic attractive interaction between the antigen HR2 peptide and the heavy chain of the optimized antibodies.

### 2.10 Trajectory Tracing Reveals Walks Across Rugged Fitness Landscapes

We visualized the trajectory of the antibodies produced by our methods on a t-SNE plot starting from its seed antibody Ab-14. In the plots in Figure 8, we colored the points according to their affinity values 1*/K*_*D*_ where paler yellow color is higher affinity. The t-SNE plot revealed local clusters with high fitness values localized to specific regions, suggesting that certain sequence mutations consistently yield superior fitness. Additionally, a smooth gradient in the affinity landscape is observed from right to left, transitioning from lower to higher predicted affinity. Figure 8A shows the evolution trajectory following the Simulated Annealing (SA) approach with ABModel substitution probabilities. While most of these points were the product of single mutations, in some cases the algorithm chose a candidate of lower affinity to explore further, which allowed the method to escape local optima to progress to the global optimum. Thus, despite sometimes admitting lesser affinity values than were generated in the previous steps, the algorithm explored novel sequence space neighborhoods while compromising the fitness function. Figure 8B shows the evolutionary trajectory following the Genetic Algorithm (GA) approach, again with ABModel substitution probabilities. In each generation, GA explores some improved binding sequences as well as some poorer binding sequences, and selects a combination of stronger and weaker binding sequences for the next generation. Therefore, we can observe more weak-binding antibodies in the landscape explored by GA than by SA. Moreover, GA handles a population of antibodies in each generation and the best among them is considered the “generation best”. It always moves toward the best solution in each generation. We can observe a reflection of this property in the evolutionary trajectory in Figure 8B.

**Figure 8:**
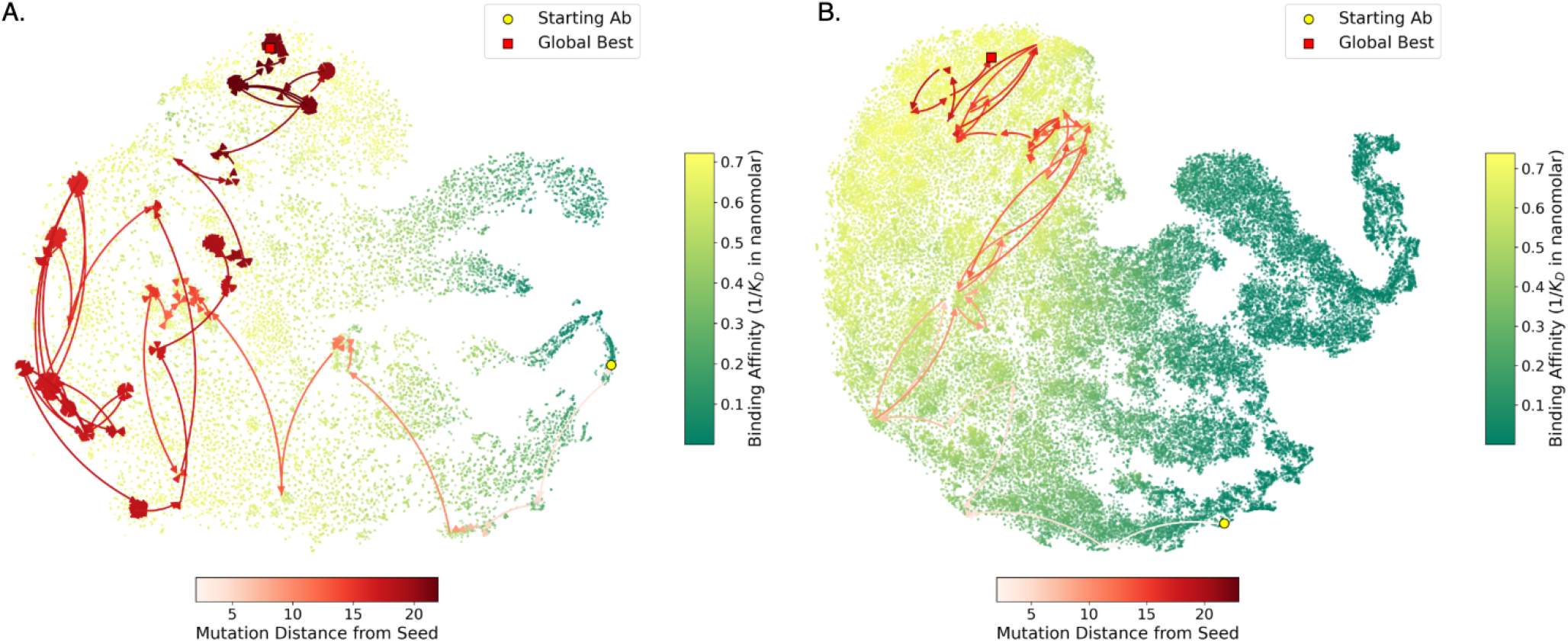
Exploring the “fitness landscape” of antibodies to find the best binding antibody; t-SNE plots of the antibody sequences generated by two methods are overlayed with the fitness function (here, the reciprocal of *K*_*D*_); variant generation and optimization were by Simulated Annealing (SA) with the ABModel substitution probabilities (panel **A**), and by Genetic Algorithm (GA) with ABModel substitution probabilities (panel **B**); the intensity of color of the arrows is mapped to the mutational distance from the respective seed (circle); globally optimized candidate antibody is labeled (box) in each panel; values of the affinity constant are mapped to green-yellow color shades.

### 2.11 Ab-Affinity’s Synthetic Antibodies Recapitulate Properties of Natural Antibodies

We compared the sequence similarity among the CDRs of our synthetic antibodies with CDRs of all known antibodies present in the SAbDab database, available through the Protein Data Bank [56]. The Jensen-Shannon divergence between these two distributions was 0.187, and the Hellinger distance was 0.191, suggesting that the distributions are comparable (**Supplementary Figure 17**). The residue distributions were indistinguishable except for those for alanine (A) and lysine (K). **Supplementary Table 1** lists the top twenty hits from the SAbDab database. Our synthetic antibodies have a sequence identity to their natural counterparts higher than 71%, especially antibodies targeting Sarbecovirus antigens.

## 3 Discussion

### 3.1 Ab-Affinity Outperforms Existing Language Models

We have developed Ab-Affinity, a large language model that can predict the binding affinity of antibodies against the SARS-CoV-2 spike protein. Ab-Affinity demonstrated stronger predictive performance than previously published methods, namely DG-Affinity, ESM-2, and AbLang. The t-SNE visualization of Ab-Affinity’s embeddings suggested that it captured meaningful relationships between antibody sequences and binding affinities to the target antigen. Unlike existing approaches that focus on generalized models for different antibody-antigen complexes (that is, DG-Affinity [14]), Ab-Affinity is tailored to capture the subtle effects of CDR mutations on their binding affinities against the HR2 segment of the SARS-Cov-2 spike protein. This capability is important for addressing the specificity of the antibody, a challenge that conventional models often fail to address. This is especially important in predicting binding affinities for mutated antibodies from a seed antibody. The superior performance of Ab-Affinity over ESM-2 and AbLang is likely because the latter are trained on “general” protein sequences. Unlike these earlier models, which are designed primarily to extract features represented by amino acid sequences, Ab-Affinity is purpose-built to predict binding affinities.

Ab-Affinity’s embeddings enabled a strong performance on several classification tasks (binding affinity, and binding improvement prediction), emphasizing the utility of our model beyond just affinity prediction. Since Ab-Affinity is fine-tuned from ESM-2, a pre-trained protein language model that represents protein sequences, Ab-Affinity inherently retains this capability. The fine-tuning process further equips Ab-Affinity to learn the effects of mutations.

The residue-residue attention maps extracted from Ab-Affinity reveal that our model focuses on the CDRs, especially CDR3, or the immediately neighboring regions. The disparities in the residue-residue attention maps highlight Ab-Affinity’s proficiency in distinguishing between strong and weak binders. Surprisingly, the model appears to have learned subtle dynamical rules of interaction between flexible epitope and paratopes, and produced predictions of intricate stabilizing interactions which were not explicitly presented during the training process. The model’s self-attention mechanism inherently learns to identify regions within the antibody that influence binding characteristics. Although CDRs are the most attended regions, the model also assigns attention to select positions outside the CDRs, reflecting its capacity to capture the effects of mutations across the entire sequence. This observation suggests that the model may inform antibody design by learning and implementing critical high-dimensional and dynamical molecular interaction rules that govern binding characteristics. In fact, despite validating similar trends, some discrepancies observed in comparisons with molecular simulations underscore the ability of Ab-Affinity to more effectively internalize the rules of molecular dynamics than the simulations themselves. Canonical molecular simulations typically capture thermal fluctuations within the observation period of the trajectories. Notably, deviations tend to increase in models that rely on rigid structures, as seen in docking approaches and coarse-grained electrostatic models.

Ab-Affinity demonstrated a better “understanding” of antibody thermal stability than ESM-2 despite not being explicitly trained on this attribute. One explanation of this finding is that the fine-tuned language model has learned intrinsic rules that govern not only the physics of antigen-antibody interaction but also the physics of protein stability that contribute to the stable interactions over a range of temperatures. Since the thermal stability of proteins depends on long-range interactions and epistasis or nonadditive influences [57, 58], the embedding provided by Ab-Affinity appears to have captured these complex nonlinear interactions better than generic protein language model embeddings.

### 3.2 The Basis of Robustness of the Generated Antibodies

In our pursuit of designing “better” antibodies by introducing mutations to the seed antibodies, we explored eight generative strategies for antibody design using Ab-Affinity as the performance metric. We found that all methods, except uniform random generation, produced synthetic antibodies with significantly improved predicted binding affinities, in some cases up to 168 times better than seed antibodies. In particular, these synthetic antibodies also outperformed all antibodies in the training library. Among the methods, PSSM showed the least improvement, probably due to its reliance on amino acid frequencies from the training dataset and its conservative substitution strategy, which tends to preserve sequence similarity to the seed. In contrast, genetic algorithms and simulated annealing proved more effective by exploring the fitness landscape more broadly: GA through recombination that introduces large sequence diversity, and SA through a gradual, probabilistic exploration of local sequence variants. While PSSM promotes conservative changes, and uniform random introduces excessive variability, methods like AbModel strike a balance by leveraging substitution matrices aligned with known antibody amino acid distributions. This balance enables the generation of novel yet biologically plausible antibodies, as supported by their consistency with established substitution probabilities. These findings highlight the importance of tailoring mutation strategies not just for novelty, but also for compatibility with biologically relevant sequence constraints. Our molecular dynamics simulations further validated the stability and specificity of the antibody-antigen complexes. The molecular dynamics analysis revealed that the synthetic antibodies are stable and form, on average, more hydrogen bonds in their interactions with the SARS-CoV-2 spike protein HR2 peptide, the target epitope.

The analysis across Ab-14, Ab-91, and Ab-95 demonstrates that GA and SA methods, especially when combined with PSSM and ABModel, significantly improve binding characteristics. This suggests that some antibodies that are more distant from the seed have better binding affinity. However, while exploring the landscape, it is critical to move toward regions with better binding affinities. Both GA and SA incorporate mechanisms to balance exploration and exploitation by selecting weak-binding antibodies and further refining them, ensuring that the generative process consistently progresses toward regions with improved binding antibodies. However, this was not always the case, as the optimization appeared to have traveled occasionally to less fit regions of the fitness landscape. It will be interesting to see whether natural variations at V(D)J junctions that encode the CDR sequences also undergo such retrograde steps during natural affinity maturation during adaptive immunity. The selection coefficient *ω* that measures the selection pressure on the CDR affinity maturation by somatic hyperrecombination can be as high as 0.9 [59]. However, they are nearly all amino acid substitutions [59], recapitulating at least one aspect of the variant generation processes adopted here. However, the estimates of *ω* are based on bulk sequence data, not single-cell lineage studies. Due to the high estimated selection coefficient, antibody-producing B cells are generally thought to be selectively allowed to survive and proliferate only along the direction of higher affinity. In reality, the natural affinity maturation processes might well utilize stochastic fluctuations in fitness values to jump over fitness barriers, as observed here with the generated antibodies.

Despite the remarkable advances in antibody design enabled by LLMs and diffusion models, the challenge of computationally generating antibodies capable of neutralizing pathogens remains difficult. Given the propensity of SARS-CoV-2 and other viruses to constantly mutate, late-evolving strains might prove to be more adept at evading vaccine- or previous infection-induced antibodies, and could even be more virulent. It is therefore crucial to predict the trajectory of the virus’ evolution and to design and rapidly develop antibody therapeutics to effectively neutralize novel strains of the virus.

## 4 Materials and Methods

We introduce a large language model called Ab-Affinity that can predict the binding affinities for complete single-chain variable sequences (both heavy and light). We show that the predictive ability of Ab-Affinity can be leveraged to generate *de novo* antibody sequences from a seed antibody. We investigate three approaches for the generation of new antibodies, namely (i) a Position Specific Score Matrix (PSSM) for directed evolution, (ii) a Genetic Algorithm (GA) that incorporates mutation and crossover operations, and (iii) a Simulated Annealing (SA) approach that explores sequence space through mutation operations such as PSSM-based substitution, random substitution, and an antibody-specific substitution matrix based on the ABModel [47]. We analyze the antibodies produced by methods (i-iii) that have the highest predicted affinity by comparing them to the “baseline” generative model that uses uniform probabilities, and by carrying out protein docking and molecular dynamics simulations.

### 4.1 Dataset

Ab-Affinity was trained on a dataset of single-chain fragment variable (scFv) antibody sequences with associated binding scores against a peptide in the SARS-CoV-2 HR2 region [60]. Whereas many regions of the SARS-CoV-2 spike protein have mutated significantly from the initial Wuhan strain, forming the Alpha, Delta, Gamma, Omicron, and numerous other minor variants, the chosen HR2 peptide is conserved across most variants of SARS and MERS spike proteins. The HR2 target epitope therefore is crucial for evaluating antibodies against the broader coronavirus group [60]. The dataset was obtained by introducing one, two, and three amino acid changes into the sequences of three candidate antibodies obtained from a phage display library, which bound to the HR2 antigen polypeptide, namely Ab-14-VH and Ab-14-VL, Ab-91-VH, Ab-95-VH, and Ab-95-VL [60]. The binding affinity (*K*_*D*_) of each of the 104,972 resulting sequence variants of the selected target peptide was experimentally estimated in a yeast mating assay, where mating efficiency was nominally related to the epitope-paratope binding affinity [60]. The equilibrium dissociation constant *K*_*D*_ value was estimated by this indirect competitive mating assay. In this assay, a population of antibodies enables the mating between two yeast strains, one of which displayed the antibody and the other displayed the antigen on the cell surface. The extent to which the two yeast strains expressing the binding proteins could mate is assumed to be proportional to (although not identical to) the *K*_*D*_ or the binding affinity between the antibody and the antigen. In [60], the proportions of the mated population were estimated by counting the proportion of DNA sequences of the corresponding antibody genes, and the relative sequence enrichment values were compared against a set of interacting protein pairs (internally controlled) with known *K*_*D*_ values measured directly by protein bi-layer interferometry. This provided the estimated *K*_*D*_ values in units of nanomolar concentration of each interaction by regression and extrapolation.

The experimental assay in [60] was conducted in triplicate for each antibody, thus the dataset contained three *K*_*D*_ values for each of 314,925 antigen-antibody pairs. Of these, 104,972 corresponded to unique antibodies. We preprocessed the interaction data by taking the arithmetic mean of the two closest *K*_*D*_ values for each antigen-antibody interaction and by eliminating the third value to reduce outliers. Antibodies that had missing values across all three replicates (33,138) were discarded from the analysis. After this preprocessing, the data from the remaining 71,834 antibodies was utilized for model training. **Supplementary Figure 1B** shows the number of antibodies for each of the three seeds, while **Supplementary Figure 1C** illustrates the number of antibodies associated with a given number of mutations from the corresponding seed antibodies. Ab-Affinity was trained on all antibody sequences (from each seed antibody) and their corresponding log-transformed binding affinity log_10_ *K*_*D*_. **Supplementary Figure 1A** shows the distributions of the log_10_ *K*_*D*_ values for the training antibodies.

### 4.2 Model Architecture

The architecture of Ab-Affinity is based on BERT [61], as implemented in ESM-2 [13] (see Figure 1D). We chose the BERT architecture because the training set consists of amino acid sequences with a few point mutations. BERT effectively captures long-range dependencies, providing meaningful representations that align with binding affinity changes caused by these mutations. In general, Ab-Affinity is composed of *N* sequential layers of encoder blocks. Each encoder block consists of multi-head attention layers [62] followed by feed-forward layers. The value of *N* determines the size of the model. We tested *N* = {6, 12, 33}, resulting in a model with 8*M*, 35*M*, and 650*M* parameters, respectively. We chose these values for *N* based on previous work, which analyzed the impact of model size on the performance of several downstream applications [13]. We used the last encoder layer output as the sequence representation (i.e., the latent space features, or *embedding*). Depending on the choices of *N* listed above, the embedding of the sequence had 320, 480, and 1280 dimensions, respectively. We added one fully connected layer to predict the binding affinity from the embedding.

### 4.3 Training

Mean squared error (MSE) was used as the loss function, and Adam was used for parameter optimization. We used 85% of the data to train the model (maintaining the same *K*_*D*_ distribution). The remaining 15% was used for model validation. We trained our model using four NVIDIA A100 (equipped with 80GB RAM) GPUs with a batch size of 128 for 100 epochs. For the amino-acid contact analysis, we used the method by Rao *et. al*. [63]. We fine-tuned the encoder layers of ESM-2, leveraging the knowledge acquired from the entire protein database. To understand the impact of pretrained protein knowledge on binding affinity prediction, we also trained an alternative model initialized with random weights. We recorded the best-performing model based on the Pearson correlation coefficient on the validation set for each setup of training. **Supplementary Figure 2** shows that the best-performing model, Ab-Affinity, used *N* = 33 layers and was fine-tuned from ESM-2.

### 4.4 t-SNE visualization of embeddings

The t-distributed Stochastic Neighbor Embedding (t-SNE) [64] was used to project the high-dimensional embeddings to 2D space. The dimensionality reduction was carried out via the Python scikit-learn package, using a perplexity of 200.

### 4.5 Generation of Synthetic Antibodies

We generated *de novo* synthetic antibody sequences by altering the CDRs of the seed antibodies Ab-14, Ab-91, and Ab-95, and using Ab-Affinity to predict the binding affinity as their fitness score. Lower *log K*_*D*_ values are considered to have higher fitness, thus *Fitness*(*s*) = 1*/f* (*s*) where *s* is the complete antibody sequence, and *f* (*s*) is the predicted binding affinity of *s* using Ab-Affinity.

We explored four strategies to generate new antibodies, namely (i) random mutations based on a uniform distribution, (ii) random mutations according to a position-specific scoring matrix, (iii) mutations introduced by a genetic algorithm, and (iv) mutations driven by a simulated annealing method.

#### 4.5.1 Baseline Uniform Model

To generate a random mutation in a protein *s* of length |*s* |, (1) we randomly selected a position *i* ∈ [1, | *s* |] to be mutated with uniform probability, then (2) we mutated the amino acid *s*[*i*] with one of the remaining 19 symbols with uniform probability. We used this *naive* approach as a baseline to compare with the generative capabilities of the other methods below.

#### 4.5.2 Position-Specific Scoring Matrix (PSSM)

To generate a random mutation in a protein *s*, given a position-specific scoring matrix (PSSM) [65] *P* of dimension | *s* | × 20, we (1) randomly selected a position *i* ∈ [1, | *s* |] with uniform probability, then (2) mutated the amino acid *s*[*i*] with one of the remaining 19 symbols using the probability distribution *P* [*i*]. The PSSM was obtained by computing the position-specific frequency of each amino acid in the training set. The pseudo-code for generating new sequences using PSSM can be found in the **Supplementary Methods**.

#### 4.5.3 Genetic Algorithm (GA)

We designed a genetic algorithm that starts from an initial population of randomly mutated seed antibody sequences. In each generation, sequences from the current set undergo crossover and mutation operations. The fitness of each sequence is evaluated using Ab-Affinity, which calculates its binding affinity log *K*_*D*_. The genetic algorithm selects the highest fitness sequences (i.e., the lowest log *K*_*D*_ values) along with a subset of lower fitness sequences, as inputs for the following generation as illustrated in Figure 1B. Three substitution strategies were employed to mutate a position, namely, (i) Random, (ii) PSSM, and (iii) ABModel. The random and the PSSM approaches are described above. The ABModel operates similarly to PSSM but employs a different substitution matrix specifically tailored for antibody sequences [47]. The genetic algorithm uses a temperature parameter that controls the diversity of the generated sequences. By tuning the temperature, we control the degree of randomness in the selection process, thereby fine-tuning the exploration of the fitness landscape. We set a population size of 50, the number of generations to 1,000, one mutation per generation, and a temperature of 5.0. To determine the sequences for the next generation, we selected the top 20% best-fit sequences, while the remaining 80% were chosen randomly from the rest. This strategy allowed us to retain the best-fitting sequences while also exploring other sequences in the fitness landscape. The **Supplementary Methods** contains the pseudo-code for the Genetic Algorithm, Crossover, and Mutation process.

#### 4.5.4 Simulated Annealing (SA)

We employed simulated annealing [66] to generate improved synthetic antibodies sampled from the binding affinity landscape. The method mutates a sequence by accepting moves that are less favorable with a small probability, thus preventing the steepest-descent algorithm from getting stuck in a local optimum. Figure 1C provides a schematic representation of this process. The algorithm starts from a seed antibody sequence. In each iteration, it generates neighboring sequences using the mutation techniques described above for the genetic algorithm. Among all the neighbors in the neighborhood, it selects the sequence with the lowest log *K*_*D*_ (greedy steepest-descent). The size of the neighborhood is defined by the allowed mutational distance. The algorithm always accepts the new neighbor if its predicted log *K*_*D*_ is lower. When the predicted value of the new neighbor is higher, it accepts it with probability 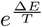 where Δ*E* is the change in predicted binding affinity and *T* is the temperature parameter. This allows the algorithm to occasionally explore uphill moves, lowering the probability that the algorithm will get stuck in a local optimum. The temperature parameter is progressively decreased by a cooling rate parameter, following the concept of annealing in physics. The algorithm continues until a maximum number of iterations is reached. In each iteration, the antibodies with the best-predicted affinity are recorded. A detailed pseudo-code of our implementation can be found in the **Supplementary Methods**.

### 4.6 Biophysical Properties

We computed the isoelectric point, hydrophobicity, instability index, and secondary structure contents for our synthetic antibodies [67]. The hydrophobicity score for each amino acid was computed using the Kyte & Doolittle index [68]. The hydrophobicity score for an antibody was obtained by computing the average hydrophobicity over all its amino acids. We also computed the (i) instability index [49] which estimates whether the antibody will be stable in solution, (ii) the fraction of alpha helices and beta sheets, and (iii) the number of peptide backbone turns in the antibodies.

### 4.7 Structural Docking

HADDOCK v2.4 [69] was used to dock the synthetic antibodies against the SARS-CoV-2 spike protein’s HR2 domain (8CZI in Protein Data Bank [56]). First, we computed the predicted 3D structure for our synthetic antibodies using ESMFold [13]. Then, we used HADDOCK to dock the predicted 3D structure with the HR2 peptide structure [70] retrieved from the Protein Data Bank (chain B). We marked the regions corresponding to the CDRs of the antibody and the peptides of the epitope so that HADDOCK could provide specific interactions between these two domains. HADDOCK score provides docking quality by combining Van der Waals, electrostatic intermolecular, desolvation, and distance restraints. We also recorded the buried surface area, which is the fraction of solvent-exposed surface area of the unbound protein that becomes solvent-inaccessible (*i*.*e*., buried) upon binding, and contributes to the Gibbs free energy of binding through entropic and enthalpic components [50].

### 4.8 Molecular Dynamics Simulation

Several combinations of seed antibodies, synthetic antibodies generated by Ab-Affinity, randomly chosen weak-binding antibodies, and the HR2 peptide structure (retrieved from the Protein Data Bank) were used for molecular dynamics (MD) simulations. All MD simulations were performed on GROMACS [71]. The models comprising the scFV-peptide complex were solvated in a cubic-water box containing 0.15 M NaCl counter-ions and its implicit TIP3P (Transferable Intermolecular Potential with Three Interaction Sites) water model [72]. The charge of the systems was neutralized by adding sodium ions. The system was minimized using a steepest-descent algorithm. The system’s pressure and temperature were adjusted to 1 atm and 300 K, in two separate 100 ps steps (NVT and NPT ensemble). The modified Berendsen and Parrinello-Rahman algorithms were applied to control the system temperature and pressure, respectively; positional restraints were applied to the protein [73, 74]. MD simulations utilized the AMBER ff99-ILDN force field [75] and the TIP3P water model [72]. The system was subjected to a 100 ns simulation at 300 K temperature, 1 bar pressure, and pH 7.0. Root Mean Square Fluctuation (RMSF) and the root mean square deviation (RMSD) of the proteins’ backbones were calculated using GROMACS tool packages. The analysis of ‘frequent conformations’ along the trajectory was performed using gmx cluster (GROMACS), based on the RMSD profile with a cutoff of 0.20 nm. The cluster with the highest population was utilized for further analysis. Generalized Born energy decomposition was performed on the last 2,500 frames with an interval of 1 with 0.15M salt by the GB-OBC2 method. The gmx MMPBSA tool [76] was employed to analyze frames corresponding to the most frequently observed clusters in each simulation run. This analysis utilizes the Poisson-Boltzmann surface area (MMPBSA) method to estimate the interaction free energies. Additionally, a per-residue energy decomposition was performed, incorporating 1-4 EEL (1-4 Electrostatic energy) into the total EEL and adding 1-4 VDW (van der Waals energy) to the overall VDW potential terms. Pymol [54] and Visual Molecular Dynamics [77] were used to visualize protein trajectories. The interacting residues were determined by MAPIYA [53] using the coordinate file from the middle frame of the most frequent cluster.

### 4.9 Constant-pH Coarse-Grained Modeling for Binding Free Energy Calculations

The fast constant-pH coarse-grained simulation method FORTE was employed to estimate the free energy of binding between the antibodies and the HR2 viral peptide from the SARS-CoV-2 spike protein [78, 79]. This biophysical model, specifically designed for protein-protein complexation studies, incorporates pH effects and focuses on electrostatic interactions [51]. The synthetic antibodies produced by Ab-Affinity were subjected to exhaustive simulations using FORTE. Two structural sets of the HR2 peptide were tested: (i) coordinates extracted from crystallographic data (chain B) and (ii) coordinates for a shorter HR2 peptide sequence (PDVDLGDISGINASVVNIQKEIDRLNEVAKNLNESLIDLQELGKYEQYIK) generated by PEP-Fold4 [80] under default parameters at pH 7 and 150 mM salt. Due to the rigid macromolecular descriptions in FORTE, the latter set was included to explore a more realistic peptide conformation in aqueous solution. However, as PEP-Fold4 imposes a limit of 50 amino acids, the five N-terminal residues, corresponding to the less structured portion of the peptide, were excluded for these tests. The core of FORTE integrates a fast proton titration scheme [81] with the capability to translate and rotate macromolecules in three dimensions using the Metropolis Monte Carlo method [78, 79]. Its key advantage lies in accurately modeling electrostatic interactions with a reduced number of sites, resulting in a smoother energy landscape. This balance offers an optimal cost-benefit ratio, enabling longer runs across multiple systems and facilitating semi-quantitative comparisons [51, 79]. In these simulations, radial distribution functions between the centers of mass of the antibody and viral peptide molecules were calculated and subsequently converted into potentials of mean force, enabling the extraction of binding affinities.

Additional runs were performed with mutants of both the antibody and the HR2 viral protein to validate the electrostatic characteristics underlying the complexation mechanism and to assess the significance of specific titratable amino acids identified in the previous analysis. The mutants were generated by neutralizing select residues from the CDRH3 region of the antibodies (ASP90, ASP106, ARG100, and LYS98) and the antigen (ASP7, ASP9, and ASP12).

### 4.10 Sequence Analysis

We computed the sequence similarity between the CDRs of synthetic antibodies and the CDRs of all known antibodies (stored in the SAbDab database [82]) by computing the Levenshtein edit distances. We also compared the frequency distributions of amino acid residues in the synthetic CDRs with those in all known CDRs. Jensen-Shannon divergence *DJS* and Hellinger distance *DH* were used to compare the distribution of amino acids in known CDRs (*P*) with the synthetically generated CDRs (*Q*). These two metrics are defined as follows

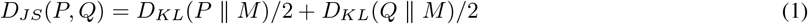

where *M* = (*P* + *Q*)/2 and 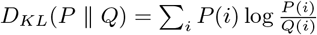, and

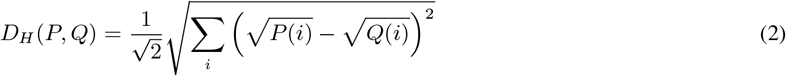

## 5 Data and Code Availability

The code, data availability, and installation instructions are available at https://github.com/ucrbioinfo/AbAffinity.

## 6 Acknowledgments

This work was funded in part by the National Institute of Allergy and Infectious Diseases (NIAID) of the US National Institutes of Health (NIH) under the award 3R01AI169543 to S.L., M.S., and A.R., and also was supported in part by the Fundação de Amparo a` Pesquisa do Estado de São Paulo (Fapesp) [Grant 2020/07158-2 to F.L.B.d.S.] and the Conselho Nacional de Desenvolvimento Científico e Tecnológico (CNPq) [Grant 305393/2020-0 to F.L.B.d.S.]. A.R. acknowledges access to the Center for Advanced Research Computing infrastructure LAGUNA HPC cluster of the University of Southern California for MD simulation through the SoCal Research Computing Alliance; F.L.B.d.S. acknowledges the Swedish National Infrastructure for Computing (SNIC) at NSC and PDC for providing resources used to run the constant-pH simulations; S.L. acknowledges the High Performance Computing Center at University of California Riverside for the training of AB-Affinity.

