## supplementary for "A Large Language Model Guides the Affinity Maturation of Antibodies Generated by Combinatorial Optimization Algorithms": Supplementary___LLM_antibody_affinity_and_generation__Copy___Copy_ (1).pdf

(Supplementary Material)

Faisal Bin Ashraf, Karen Paco, Zihao Zhang, Christian J. Dávila Ojeda, Mariana P. Mendivil, Jordan A. Lay, Tristan Y. Yang, Fernando L. Barroso da Silva, Matthew H. Sazinsky, Animesh Ray, Stefano Lonardi

#### Supplementary Methods

**Algorithm 1** PSSM

**Require:** $S$ *// list of sequences*
**Ensure:** $PSSM$, $amino\_acids$, $N$

$amino\_acids \leftarrow \{A, C, D, E, F, G, H, I, K, L, M, N, P, Q, R, S, T, V, W, Y\}$
$PSSM \leftarrow$ matrix of shape $len(S) \times len(amino\_acids)$
$l \leftarrow length\ of\ sequences\ in\ S$
**for** $i = 1 \ldots len(S)$ **do**
  **for** $j = 1 \ldots l$ **do**
    $idx \leftarrow$ index of $S[i][j]$ in $amino\_acids$
    $PSSM[j][idx] += 1$
  **end for**
**end for**
$PSSM \leftarrow \frac{PSSM}{len(S)}$ *// normalize PSSM matrix*
$S_n = \phi$ *// list for new generated sequences*
**for** $i = 1 \ldots N$ **do**
  **for** $j = 1 \ldots l$ **do**
    $Seq_i[j] \leftarrow$ select an amino acid using the probability in $P[j]$
  **end for**
  $S_n \leftarrow S_n \cup Seq_i$
**end for**
**return** $S_n$

**Algorithm 2** Genetic Algorithm

**Require:** $P_{init}, N, mutation\_rate, k$
**Ensure:** $P$

```
P ← P_init
for N times do
    C ← Crossover(P) // crossover of sequences in P_init
    M ← Mutation(C, mutation_rate) // mutation of sequences in C
    P_gen ← P ∪ C ∪ M
    P_top ← Top k sequences from P_gen using the score from Ab-Affinity
    P_low ← Remaining size(P) − k random sequences from P_gen − P_top
    P ← P_top ∪ P_low
end for
return P
```

**Algorithm 3** Crossover

**Require:** $P$

```
P_new ← ϕ
l ← length of sequences in P
for i = 1 . . . size(P) do
    c ← randomly selected point from 1 . . . l
    C_1 ← P[i][1 : c] + P[i + 1][c : l]
    C_2 ← P[i + 1][1 : c] + P[i][c : l]
    P_new ← P_new ∪ C_1 ∪ C_2
end for
return P_new
```

**Algorithm 4** Mutation

**Require:** $P, mutation\_rate$

```
P_new ← ϕ
l ← length of sequences in P
for i = 1 . . . size(P) do
    for j = 1 . . . l do
        P[i][j] ← select new amino acid with mutation_rate probability
    end for
    P_new ← P_new ∪ P[i]
end for
return P_new
```

**Algorithm 5** Simulated Annealing

**Require:** $S_{init}$, $T$, $cooling\_rate$, $N$
**Ensure:** $S_{curr}$, $E_{curr}$, $S_{best}$, $E_{best}$

$S_{curr} \leftarrow S_{init}$
$E_{curr} \leftarrow score\_sequence(S_{curr})$ *// predicted affinity of $S_c$ using Ab-Affinity*
$S_{best} \leftarrow S_c$
$E_{best} \leftarrow E_c$
**for** $i \leftarrow 1$ **to** $N$ **do**
    $S_{new} \leftarrow explore\_nearby\_sequences(S_{curr}, mutate)$ *// get mutated sequences from $S_{curr}$ using $mutate()$ function and select the best as $S_{new}$*
    $E_{new} \leftarrow score\_sequence(S_{new})$
    $\Delta E \leftarrow E_{new} - E_{curr}$
    **if** $\Delta E < 0$ **or** with $e^{(\Delta E/T)}$ probability **then**
        $S_{curr} \leftarrow S_{new}$
        $E_{curr} \leftarrow E_{new}$
    **end if**
    **if** $E_{curr} < E_{best}$ **then**
        $S_{best} \leftarrow S_{curr}$
        $E_{best} \leftarrow E_{curr}$
    **end if**
    $T \leftarrow T \times cooling\_rate$
**end for**
**return** $S_{best}, E_{best}$

**Bond Free Energy Estimation.** For hydrogen bonds, we used the approximate equation [A],

$$\Delta G_{H-bond} = \alpha \left(1 - \frac{r}{r_0}\right) \tag{1}$$

where $r$ is the observed bond distance and $r_0$ for hydrogen bonds = 2.8°A, and $\alpha$ = 2.0 kcal/mol.

The energy of interaction between two charges is estimated by Coulomb's law, as follows.

$$E_{electrostatic} = \frac{q_1 q_2}{4\pi\epsilon_0\epsilon_r r} \tag{2}$$

where $\epsilon_0$ is the permittivity of free space $(8.854187817 \times 10^{-12} C^2/Jm)$, $\epsilon_r$ is the dielectric constant of a protein (close to 10, dimensionless unit), and $r$ is the distance of separation between the two charges (in meter), $q_1$ and $q_2$ (here, +1 and -1). Thus, the total bond energy is

$$\Delta G_{total} = \Delta G_{H-bond} + E_{electrostatic} \tag{3}$$

### Supplemental Tables and Figures

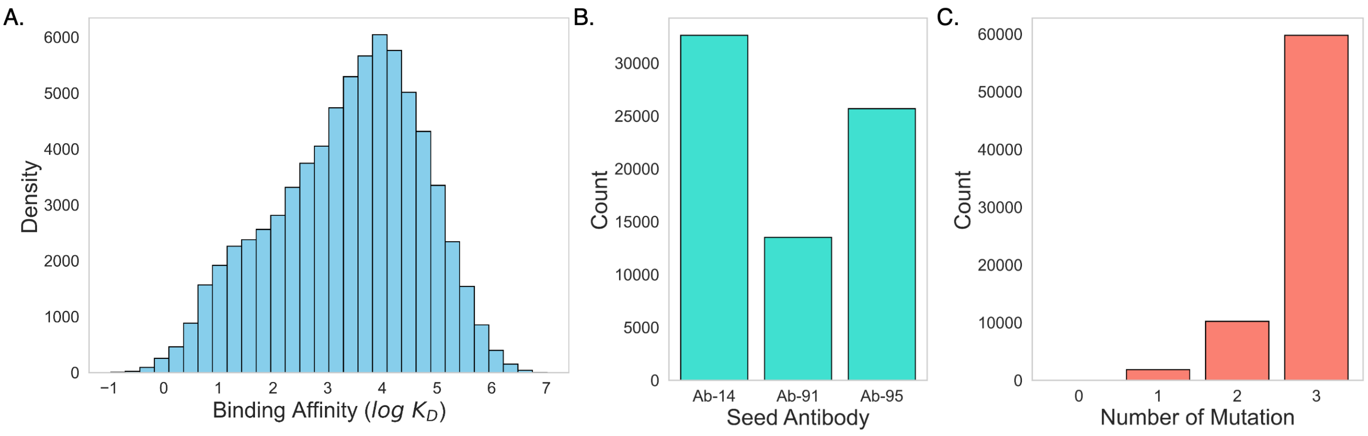

Supplementary Figure 1: Training set summary statistics. **A** Distribution of binding affinity $\log\ K_D$; **B** Number of antibodies from each seed; **C** Number of antibodies with a given number of mutation from each seed.

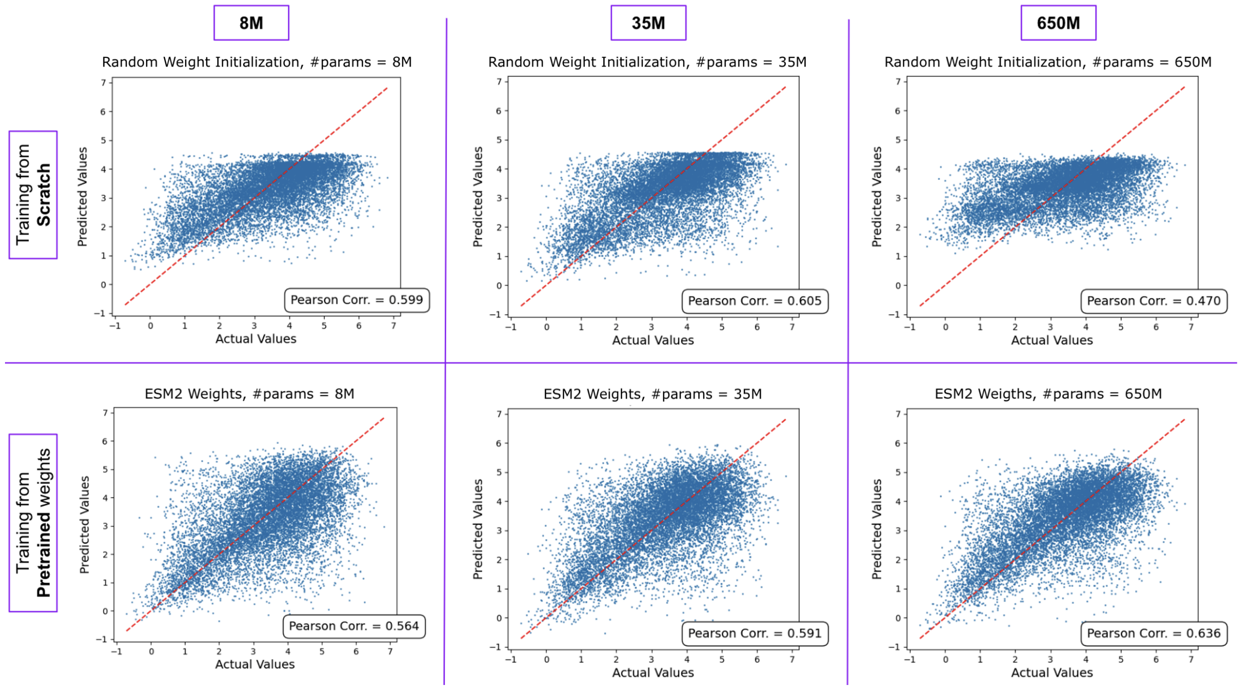

Supplementary Figure 2: Actual vs. predicted binding affinity on the test set using different predictive models. The top row represents models trained with randomly initialized weights, while the bottom row shows the performance of models fine-tuned starting from a pretrained ESM-2. The columns represent the size of the model. The leftmost column is the smallest model with 8 million parameters, while the rightmost column is the largest model with 650 million parameters. The Pearson correlations on the validation set for each model is also reported.

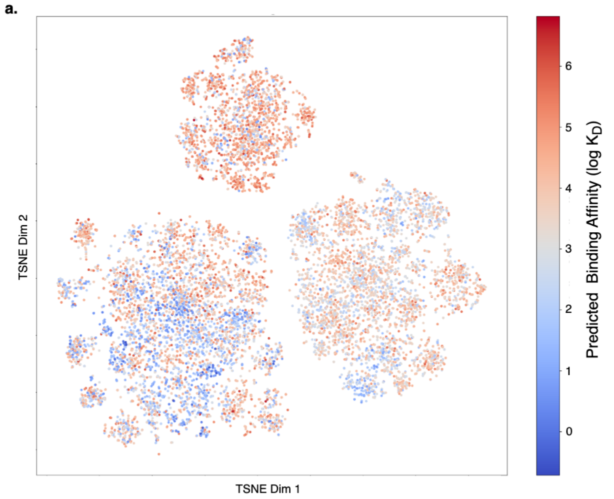

Supplementary Figure 3: t-SNE representation of the embedding produced by **(a)** ESM-2 and **(b)** Ab-Affinity; antibodies are colored according to their binding affinity.

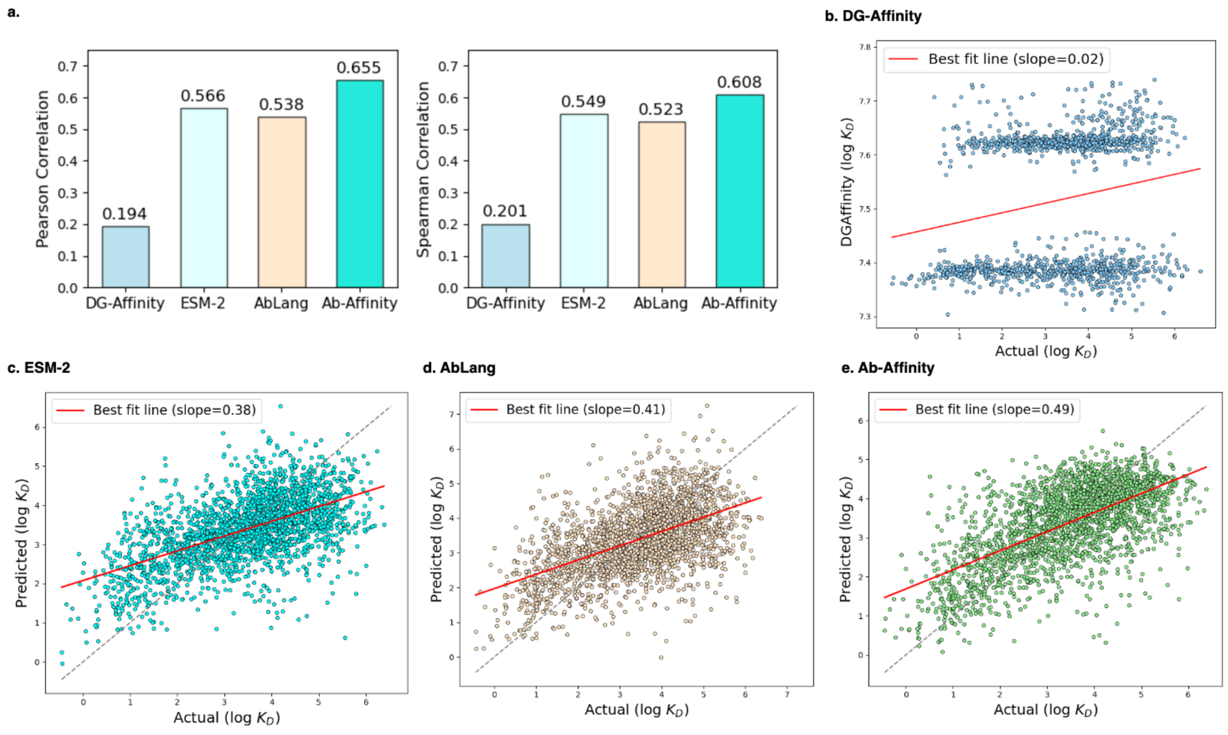

Supplementary Figure 4: Comparing affinity prediction models. **(a)** Pearson and Spearman correlation for DG-Affinity (p-values = $3.86 \times 10^{-14}$, and $3.38 \times 10^{-15}$), ESM-2 (p-values = $8.03 \times 10^{-198}$, and $8.02 \times 10^{-198}$), AbLang (p-values = $1.24 \times 10^{-175}$, and $1.02 \times 10^{-163}$) and Ab-Affinity (p-values = $4.03 \times 10^{-261}$, and $8.65 \times 10^{-217}$); scatter plot for actual vs. predicted binding affinity for **(b)** DG-Affinity, **(c)** ESM-2, **(d)** AbLang, **(e)** Ab-Affinity (includes antibodies for all three seeds).

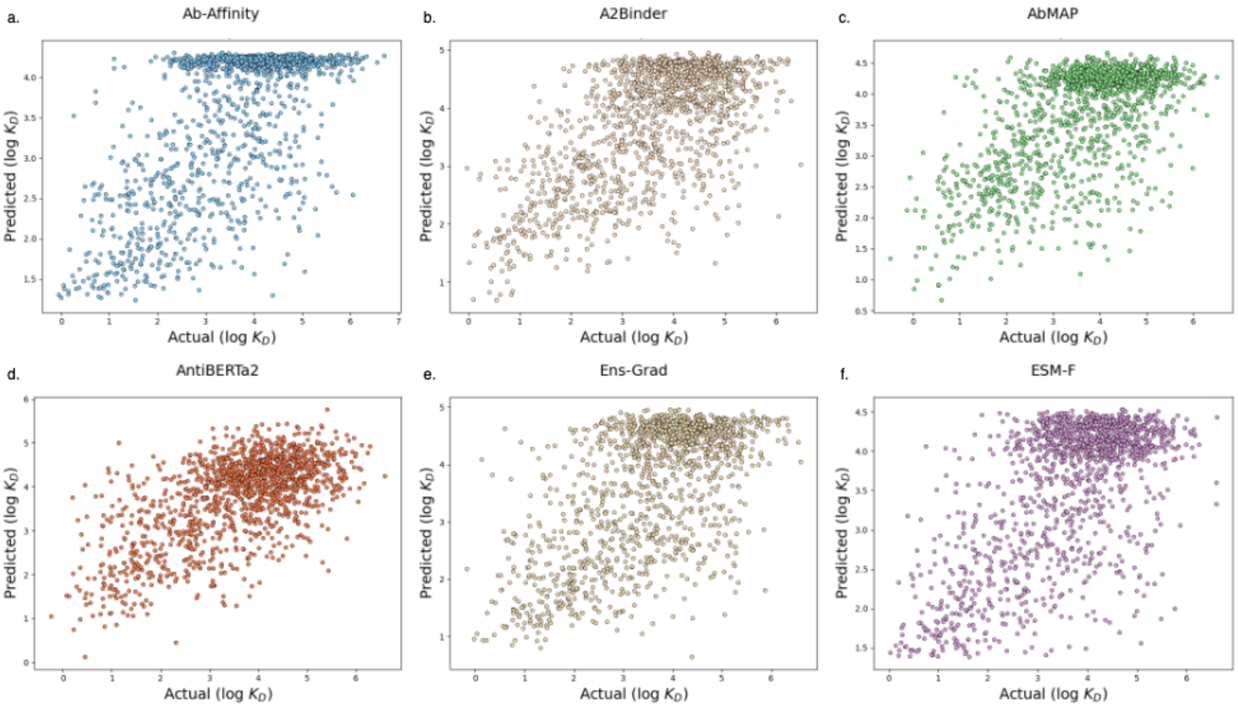

Supplementary Figure 5: Actual vs. predicted binding affinity on the 14H dataset for various methods.

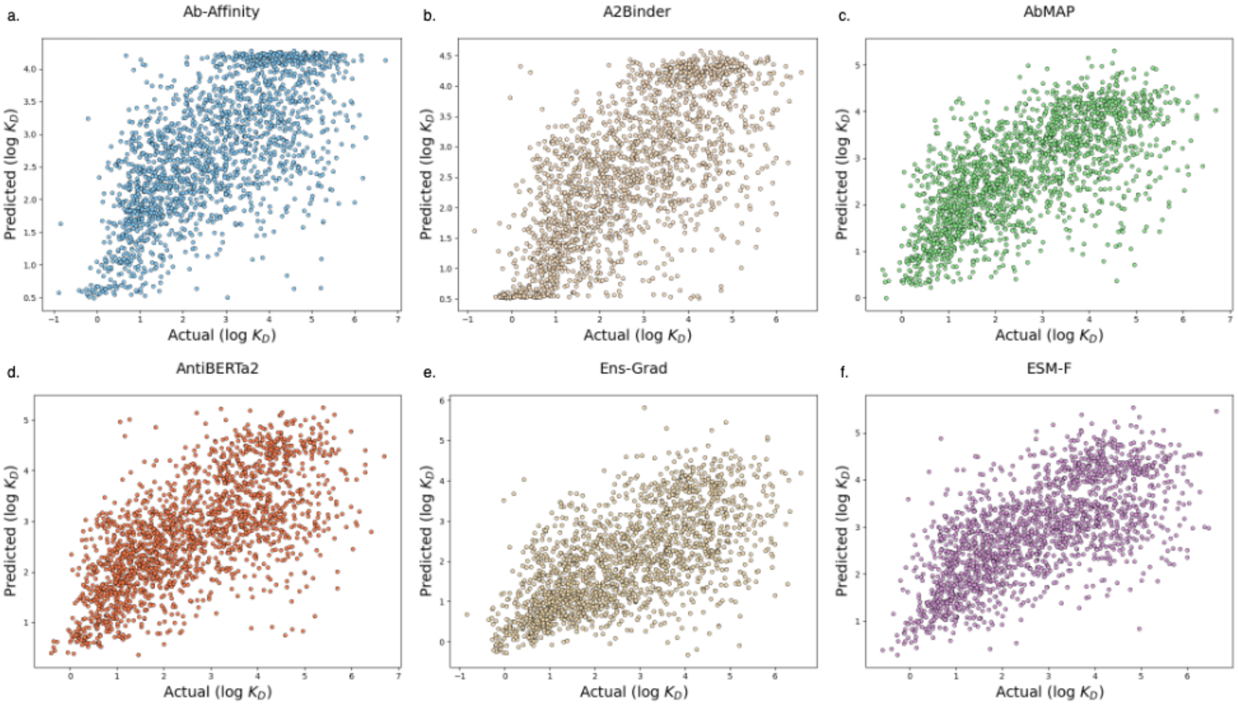

Supplementary Figure 6: Actual vs. predicted binding affinity on the 14L dataset for various methods.

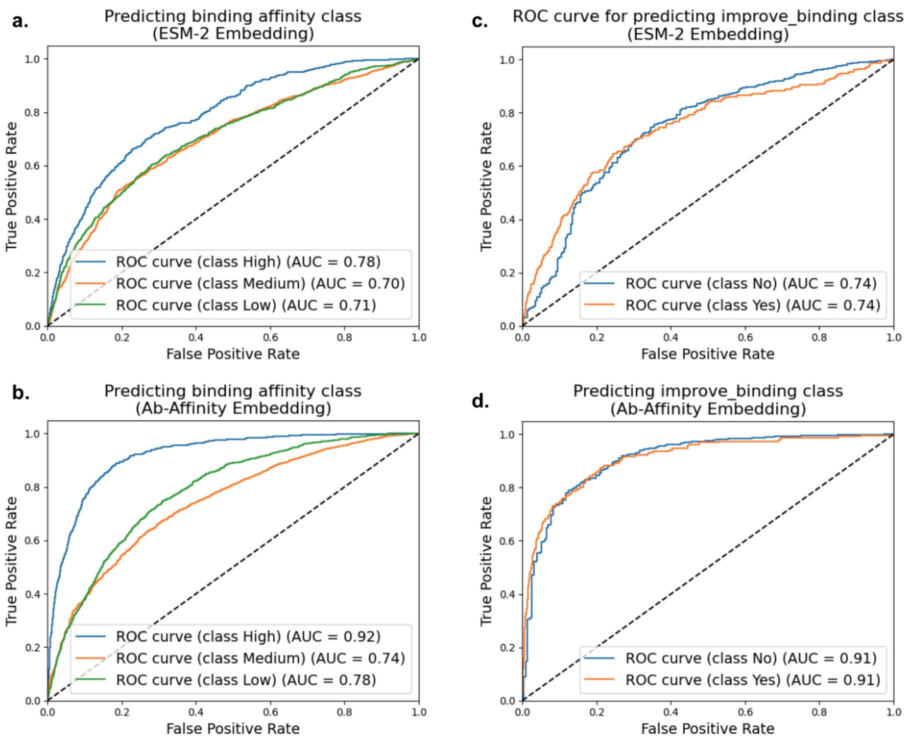

Supplementary Figure 7: ROC curves and AUC values for two classification tasks, namely **(a,b)** determining the binding affinity class of an antibody (High, Medium, Low) and **(c,d)** determining whether the binding is improved compared to the seed antibody (Y/N); **(a,c)** ROC curves using the ESM-2 embedding; **(b,d)** ROC curves using the Ab-Affinity embedding; AUC values are reported for each classifier.

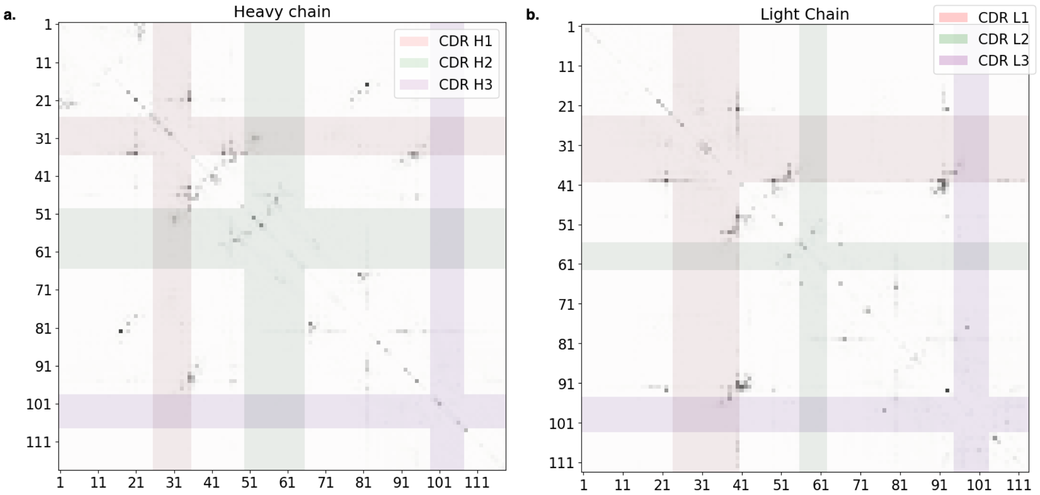

Supplementary Figure 8: Differences between the residue-residue attention maps for strong binding (i.e. $\log K_D < 0.5$) and weak binding (i.e., $\log K_D > 5.5$) for antibodies generated from Ab-14 **(a)** heavy chain, and **(b)** light chain; colored stripes highlight the CDR region; all $K_D$ values are in units of nM.

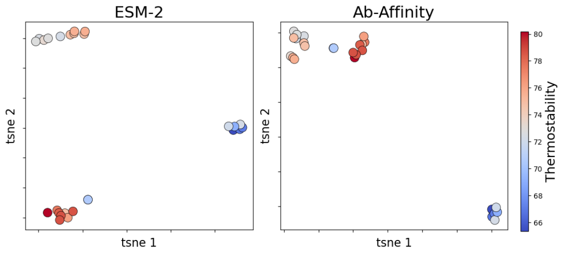

Supplementary Figure 9: t-SNE visualization of the embeddings produced by ESM-2 and Ab-Affinity; points are colored according to the experimentally-determined thermostability of those antibodies.

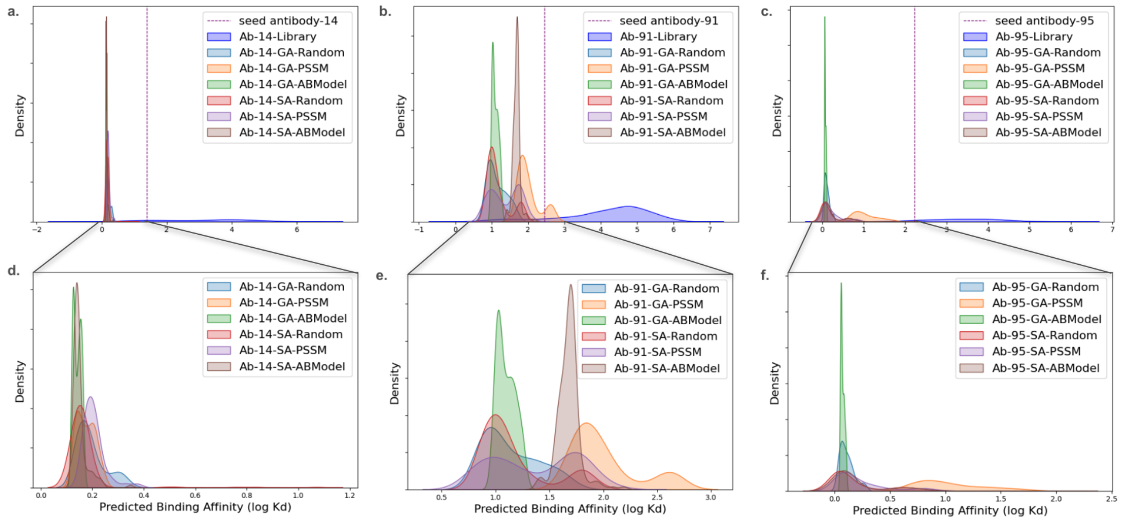

Supplementary Figure 10: Distribution of predicted binding affinity for the antibodies in the training library compared to the synthetic antibodies generated by the GA and SA methods: (a,d) Ab-14, (b,e) Ab-91 and (c,f) Ab-95

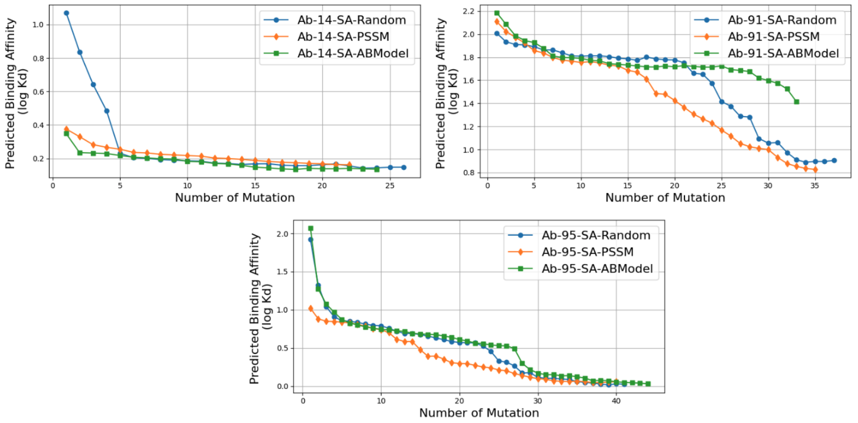

Supplementary Figure 11: Predicted binding affinity of the synthetic antibodies as a function of the number of mutations from the corresponding seed antibody

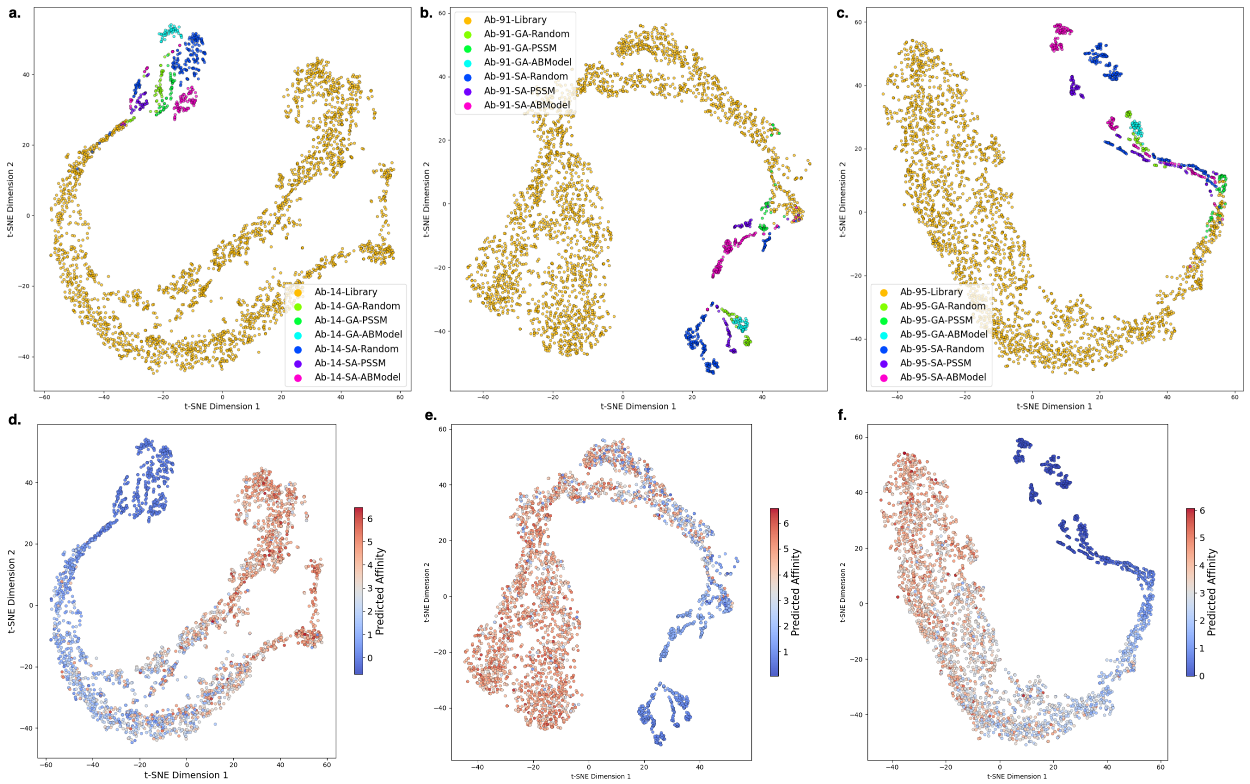

Supplementary Figure 12: t-SNE projection for the Ab-affinity embedding of the antibodies in the training library and the synthetic antibodies; in (a-c) points are colored by the method that produced it [(a) Ab-14, (b) Ab-91, (c) Ab-95]; in (d-f) points are colored by the predicted binding affinity [(d) Ab-14, (e) Ab-91, (f) Ab-95]

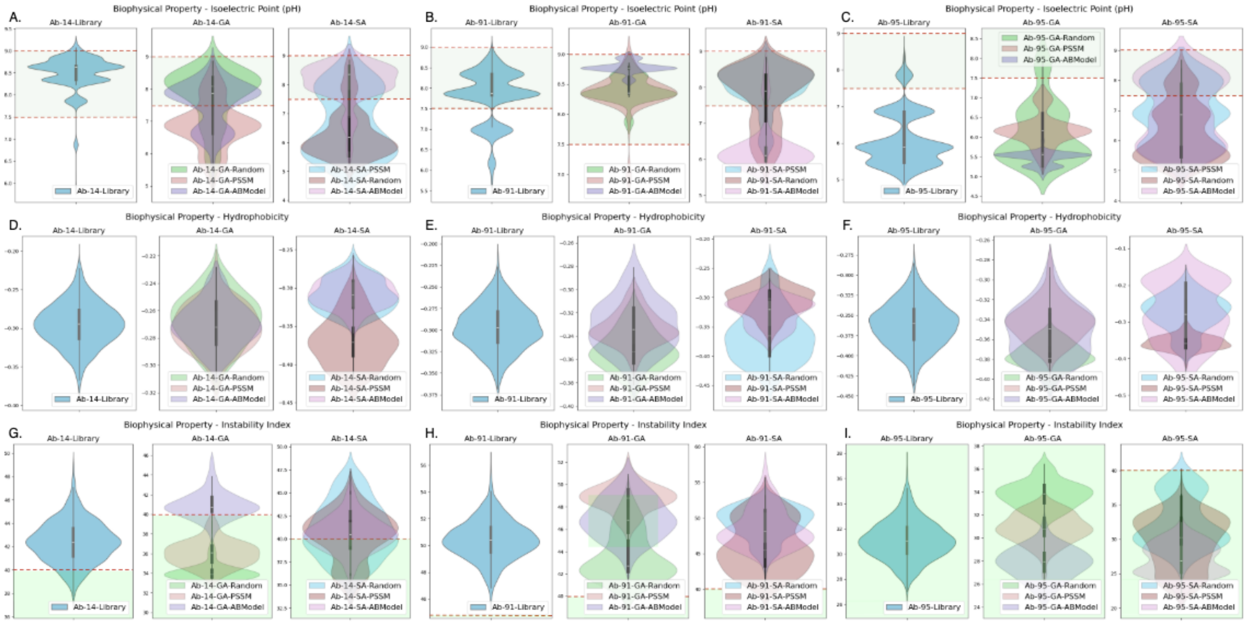

Supplementary Figure 13: Distribution of some biophysical properties for the antibodies in the training library and the synthetic antibodies. (A, B, C) distributions of isoelectric point [ideal range 7.5-9 pH]; (D, E, F) distribution of mean hydrophobicity [lower is better]; (G, H, I) distribution of instability index [less than 40 is stable]

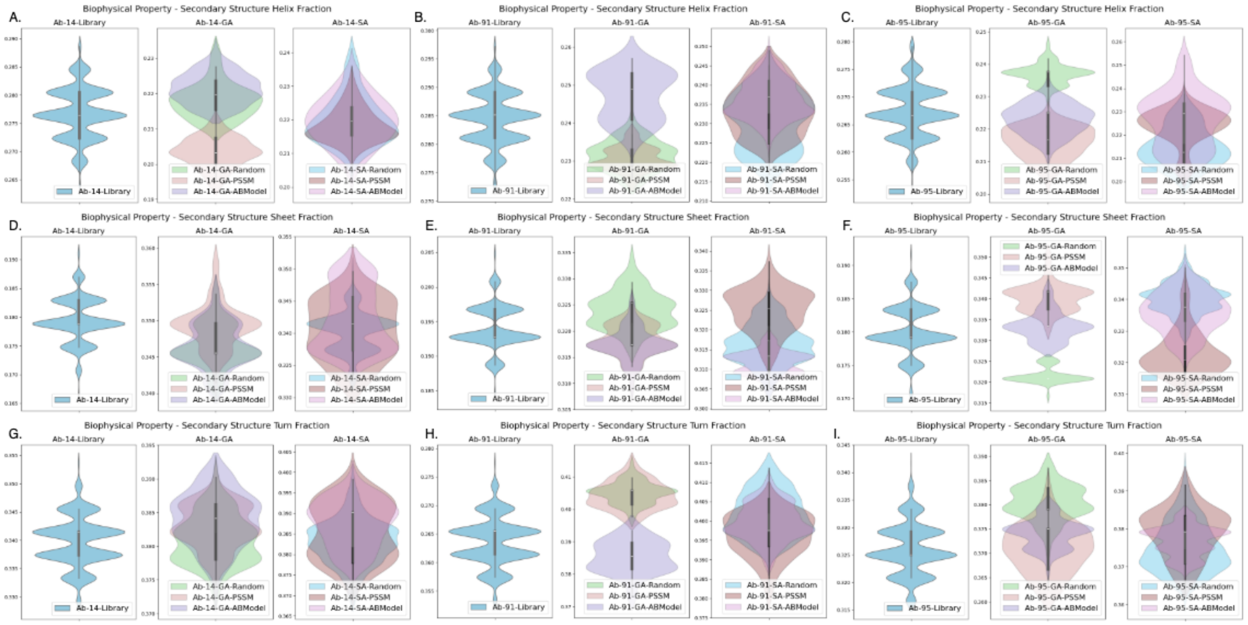

Supplementary Figure 14: Distribution of secondary structure properties for the antibodies in the training library and the synthetic antibodies. (A, B, C) Alpha Helix Fraction; (D, E, F) Beta Sheet Fraction; (G, H, I) Turn fraction

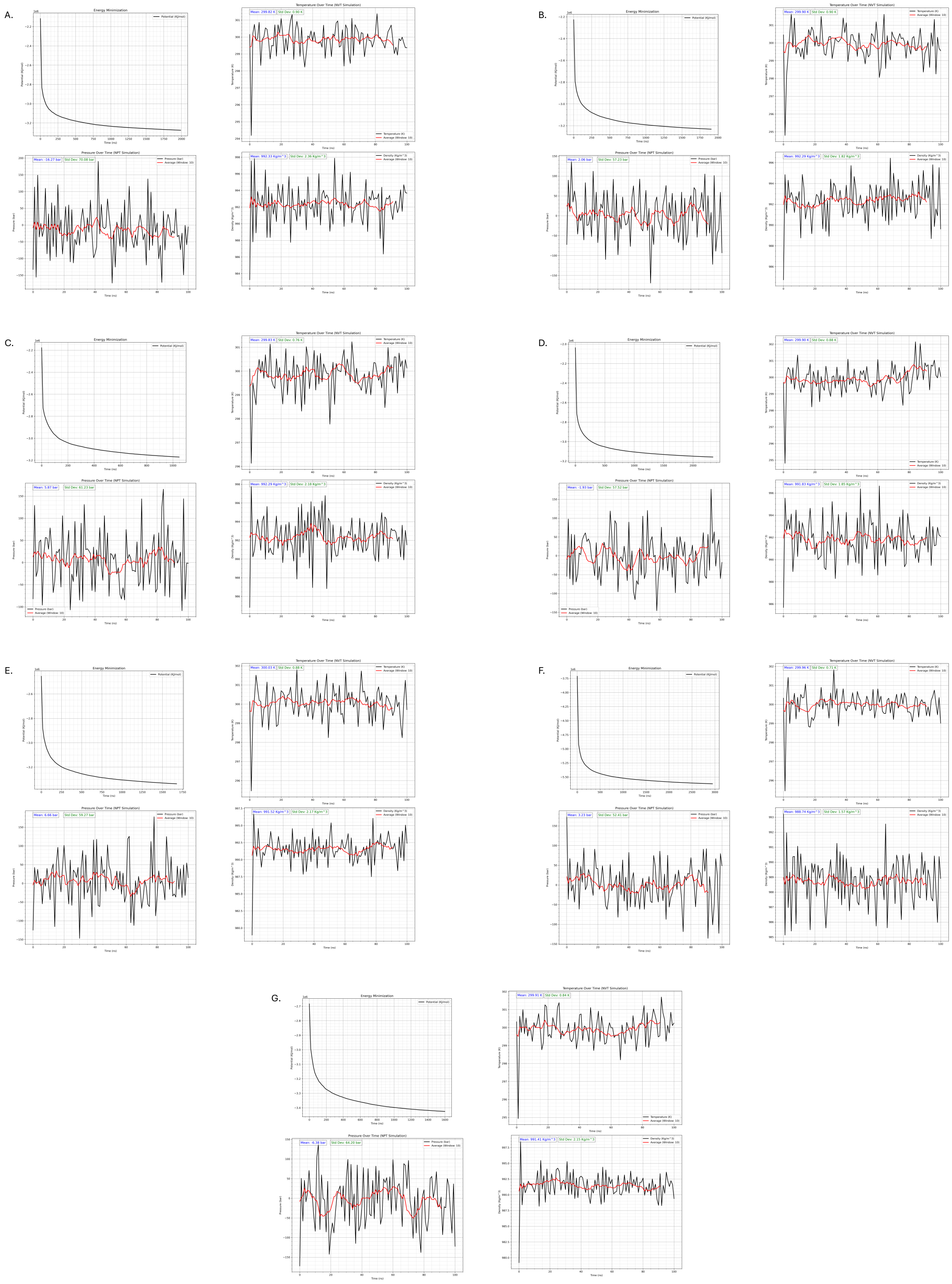

Supplementary Figure 15: Quality control plots from MD simulations of seven antibodies, Ab-14-SA-pssm-1 (A), Ab-14-SA-pssm-6 (B), Ab-14-seed (C), Ab-95-GA-random-1 (D), Ab-95-GA-random-3 (E), Ab-95-GA-random-7 (F), Ab-95-seed (G)

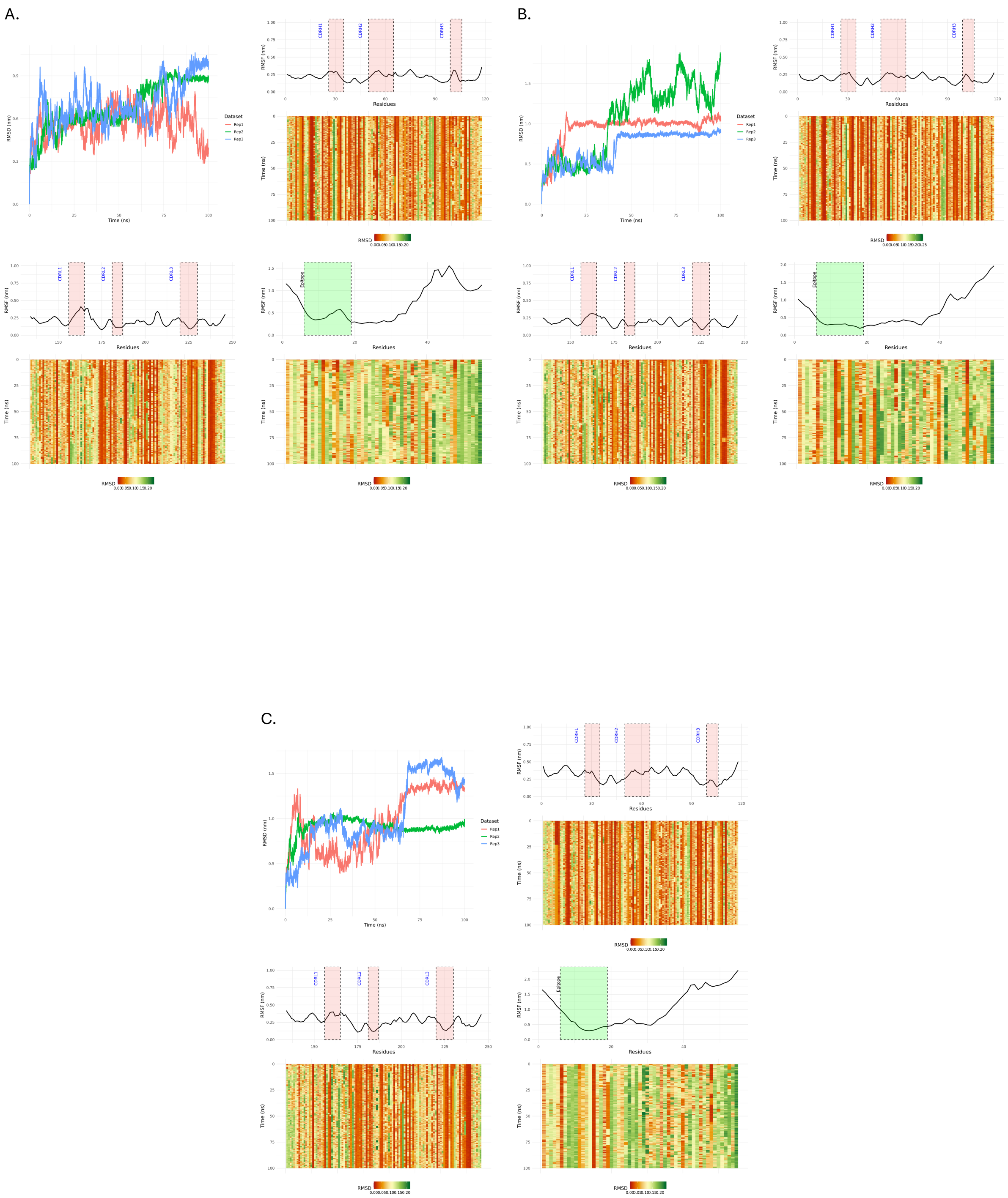

Supplementary Figure 16: Structural fluctuations observed during molecular dynamics simulations, depicted through RMSD of the backbone vs. time plots, RMSF per residue profiles, and RMSD per residue vs. time heatmaps for the heavy chain, light chain, and antigen. Results are presented for antibodies Ab-14-SA-pssm-1 (A), Ab-14-SA-pssm-6 (B), and Ab-14-seed (C). Different colors in the RMSD plots indicate distinct replicate runs.

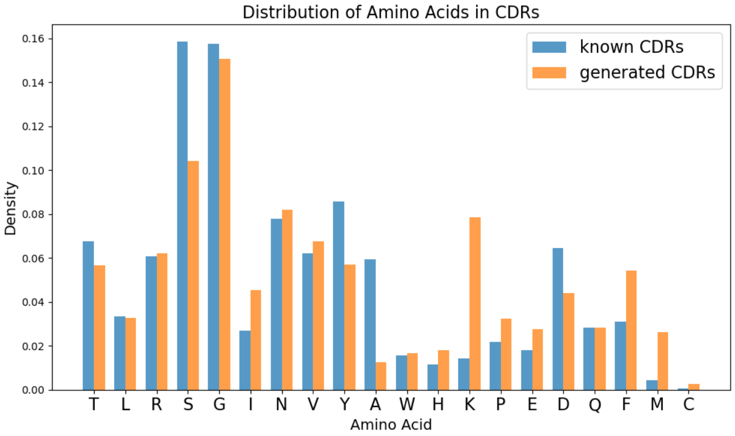

Supplementary Figure 17: Distribution of amino acids in synthetic vs. known CDRs

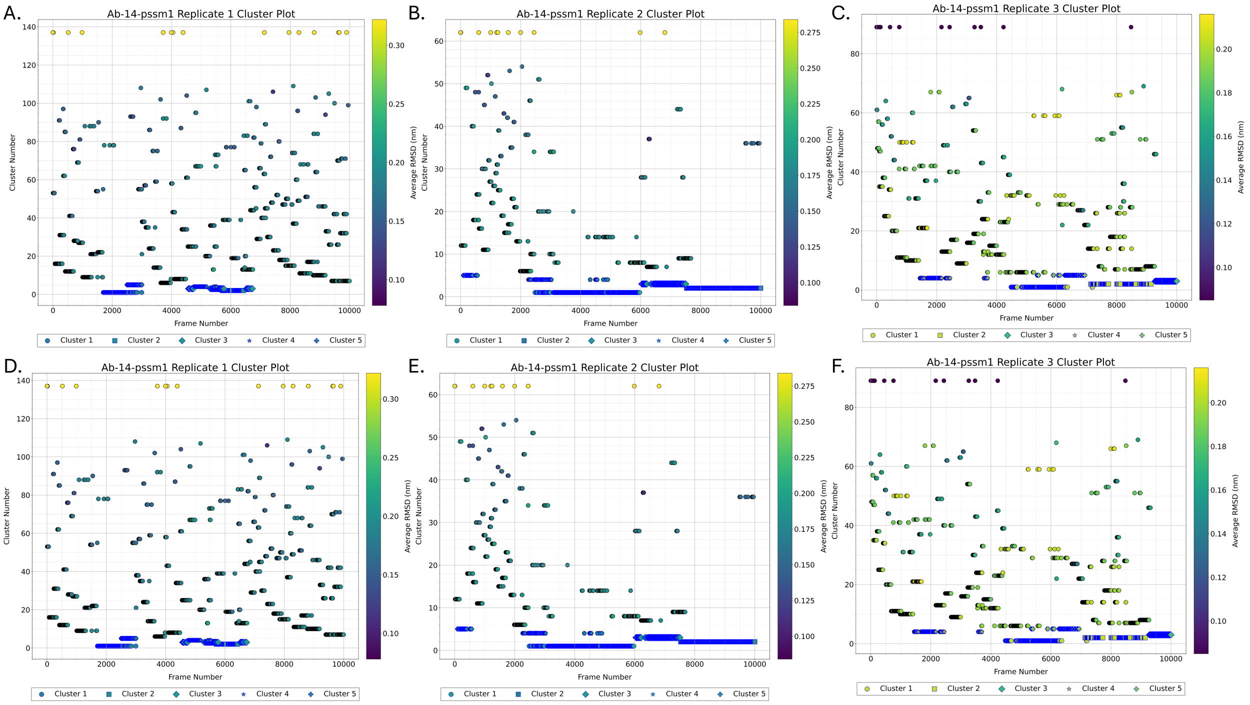

Supplementary Figure 18: Cluster analysis results from the MD trajectories obtained across three replicate runs. (a) Results from replicate runs A, B, and C for Ab-14-SA-pssm-1. (b) Results from replicate runs C, D, and E for Ab-14-SA-pssm-6.

Supplementary Table 1: Top-20 hits from the SAbDab database when queried with the CDRs of our synthetic antibodies; amino acids in red indicate the mismatches

| Seed Antibody | Synthetic CDR | Known CDR | PDB ID | Antigen | Identity |
|---|---|---|---|---|---|
| Ab-91 | TLRSGINVGTYRIY | TLRSGINVGTYRIY | 6w7s | *Saccharopolyspora Erythraea* | 1.00 |
| Ab-91 | MIWHSSAWV | AIWHSSAWV | 6kva | Peptide from CXCR2 | 0.89 |
| Ab-95 | GKNYRPA | GKNYRPS | 6fg2 | Multiple Sclerosis (MS) disease | 0.86 |
| Ab-14 | GASQRES | GASTRES | 7jln | HIV | 0.86 |
| Ab-14 | DATTRAS | DASTRAS | 7s8h | Lassa Virus | 0.86 |
| Ab-14 | DADTRAS | DASTRAS | 7s8h | Lassa Virus | 0.86 |
| Ab-14 | GASQRET | GASTRET | 8guz | Hemophilia | 0.86 |
| Ab-14 | DATTRAA | DASTRAA | 8sxj | HIV | 0.86 |
| Ab-91 | YKSDSDKQQGSGV | YKSDSDKQQGS | 6kva | Peptide from CXCR2 | 0.85 |
| Ab-95 | QGDSLRGGYAN | QGDSLRGYYAS | 6q1j | H1 influenza virus | 0.82 |
| Ab-14 | QQYHTLPLS | QQYHTTPLT | 8t7a | Respiratory Syncytial Virus | 0.78 |
| Ab-14 | QQYSRLPLS | QQYSRLPFT | 1emt | Synthetic | 0.78 |
| Ab-91 | QVWHSSAVV | QVWDSSSVV | 4jzn | Hepatitis C Virus | 0.78 |
| Ab-95 | TSRDSSGFQVF | TSRDSSGNHVI | 8khc | SARS CoV-2 | 0.73 |
| Ab-95 | QGQSLRYNYAN | QGDSLRSNYAS | 6fg2 | Multiple Sclerosis (MS) disease | 0.73 |
| Ab-95 | QGESLRYNYAN | QGDSLRSNYAS | 6fg2 | Multiple Sclerosis (MS) disease | 0.73 |
| Ab-95 | QGDSLKGGYAN | QGDSLRGYYAS | 6q1j | H1 influenza virus | 0.73 |
| Ab-95 | SSRDGSGTSVY | SSYDGSSTSVV | 1jvk | Amyloidosis | 0.73 |
| Ab-95 | GSNNRPA | GKNNRPS | 8khc | SARS CoV-2 | 0.71 |
| Ab-95 | GKSNRPQ | GKNNRPS | 8khc | SARS CoV-2 | 0.71 |

Supplementary Table 4: Statistical analysis of difference between the variances of residue RMSD distributions of candidate antibody-epitope and seed antibody-epitope complexes

| Name of Antibody | Distribution of variance in RMSD (70-100 ns) | Variance compared with RMSD of seed Ab (70-100 ns) | Levene Statistic | Levene p-value | Fligner Statistic | Fligner p-value |
|---|---|---|---|---|---|---|
| Ab-95-GA-random-3 | CDR_RMSD | Ab-95 seed CDR | 41.4569439 | 1.40E-10 | 38.59429949 | 5.22E-10 |
| Ab-95-GA-random-3 | Epitope_RMSD | Ab-95 seed Epitope | 209.2697628 | 6.72E-46 | 216.1642413 | 6.21E-49 |
| Ab-14-SA-pssm-6 | CDR_RMSD | Ab-14 seed CDR | 22.8794406 | 1.81E-06 | 23.95714979 | 9.85E-07 |
| Ab-14-SA-pssm-6 | Epitope_RMSD | Ab-14 seed Epitope | 252.3842401 | 1.25E-54 | 232.3279694 | 1.85E-52 |
| Ab-95-GA-random-1 | CDR_RMSD | Ab-95 seed CDR | 320.078232 | 4.28E-68 | 277.475721 | 2.66E-62 |
| Ab-95-GA-random-1 | Epitope_RMSD | Ab-95 seed Epitope | 109.0336959 | 4.28E-25 | 118.6542641 | 1.25E-27 |
| Ab-95-GA-random-4 | CDR_RMSD | Ab-95 seed CDR | 149.7326711 | 1.24E-33 | 136.7006297 | 1.40E-31 |
| Ab-95-GA-random-4 | Epitope_RMSD | Ab-95 seed Epitope | 1211.094563 | 3.51E-223 | 984.8529253 | 3.52E-216 |
| Ab-14-GA-random-10 | CDR_RMSD | Ab-14 seed CDR | 46.01371686 | 1.41E-11 | 45.38463161 | 1.62E-11 |
| Ab-14-GA-random-10 | Epitope_RMSD | Ab-14 seed Epitope | 49.87783809 | 2.03E-12 | 42.26228009 | 7.98E-11 |
| Ab-14-SA-pssm-1 | CDR_RMSD | Ab-14 seed CDR | 27.9135007 | 1.36E-07 | 24.76204321 | 6.49E-07 |
| Ab-14-SA-pssm-1 | Epitope_RMSD | Ab-14 seed Epitope | 2.090636163 | 0.148308871 | 1.994887909 | 0.157830742 |
| Ab-14-SA-pssm-2 | CDR_RMSD | Ab-14 seed CDR | 62.95308076 | 2.96E-15 | 68.40441994 | 1.33E-16 |
| Ab-14-SA-pssm-2 | Epitope_RMSD | Ab-14 seed Epitope | 21.81894504 | 3.13E-06 | 8.981883851 | 0.002726694 |

Supplementary Table 2: Constant-pH Coarse-Grained Modeling for Binding Free Energy Calculations. Estimated binding affinities between antibodies and HR2 peptides, expressed in $k_B$T units. Data were obtained from FORTE simulations conducted at pH 7, 150 mM NaCl, and 298 K. The antigens, named PEPFOLD-model1, HR2 Domain, HR2 Domain chainA, and HR2 Domain chainB, correspond respectively to: (1) the short 50-monomer peptide structure produced by PEP-Fold4, (2) the homotrimer peptide structure provided by PDB ID 8CZI, (3) a single peptide from chain A of the same PDB coordinates, and (4) chain B from the same PDB. The antibody names, extended as -SA-PSSM-1, -SA-PSSM-6, -seed, -GA-random-1, -GA-random-3, and -GA-random-7, are detailed in the text, where further explanations are provided.

| Antibody | Antigen | Estimated Binding Affinity | | |
|---|---|---|---|---|
| | | Repetition1 | Repetition2 | Repetition3 |
| CoV-AAYL49_4032 | PEPFOLD-model1 | −1.1203 | −1.1190 | −1.1192 |
| CoV-AAYL49_4032 | HR2_Domain | −2.4096 | −2.4107 | −2.4195 |
| CoV-AAYL49_4032 | HR2_Domain_chainA | −1.1660 | −1.1633 | −1.1661 |
| CoV-AAYL49_4032 | HR2_Domain_chainB | −1.1767 | −1.1600 | −1.1633 |
| CoV-AAYL50_21000 | PEPFOLD-model1 | −1.0721 | −1.0704 | −1.0695 |
| CoV-AAYL50_21000 | HR2_Domain | −2.3451 | −2.3611 | −2.3522 |
| CoV-AAYL50_21000 | HR2_Domain_chainA | −1.1196 | −1.1353 | −1.1278 |
| CoV-AAYL50_21000 | HR2_Domain_chainB | −1.1243 | −1.1166 | −1.1366 |
| CoV-Ab-14-SA-PSSM-1 | PEPFOLD-model1 | −0.8383 | −0.8412 | −0.8425 |
| CoV-Ab-14-SA-PSSM-1 | HR2_Domain | −1.9810 | −1.9570 | −1.9696 |
| CoV-Ab-14-SA-PSSM-1 | HR2_Domain_chainA | −0.9072 | −0.9151 | −0.9163 |
| CoV-Ab-14-SA-PSSM-1 | HR2_Domain_chainB | −0.9211 | −0.9205 | −0.9121 |
| CoV-Ab-14-SA-PSSM-6 | PEPFOLD-model1 | −0.9529 | −0.9584 | −0.9524 |
| CoV-Ab-14-SA-PSSM-6 | HR2_Domain | −2.1893 | −2.1905 | −2.1822 |
| CoV-Ab-14-SA-PSSM-6 | HR2_Domain_chainA | −1.0236 | −1.0230 | −1.0334 |
| CoV-Ab-14-SA-PSSM-6 | HR2_Domain_chainB | −1.0281 | −1.0278 | −1.0271 |
| CoV-Ab-14-seed | PEPFOLD-model1 | −1.1157 | −1.1159 | −1.1103 |
| CoV-Ab-14-seed | HR2_Domain | −2.4174 | −2.4169 | −2.4096 |
| CoV-Ab-14-seed | HR2_Domain_chainA | −1.1506 | −1.1533 | −1.1445 |
| CoV-Ab-14-seed | HR2_Domain_chainB | −1.1458 | −1.1472 | −1.1560 |
| CoV-Ab-95-GA-random-1 | PEPFOLD-model1 | −0.9011 | −0.8988 | −0.8988 |
| CoV-Ab-95-GA-random-1 | HR2_Domain | −2.1154 | −2.1032 | −2.1058 |
| CoV-Ab-95-GA-random-1 | HR2_Domain_chainA | −0.9814 | −0.9901 | −0.9810 |
| CoV-Ab-95-GA-random-1 | HR2_Domain_chainB | −0.9781 | −0.9856 | −0.9814 |
| CoV-Ab-95-GA-random-3 | PEPFOLD-model1 | −0.9090 | −0.9185 | −0.9135 |
| CoV-Ab-95-GA-random-3 | HR2_Domain | −2.1195 | −2.1271 | −2.1217 |
| CoV-Ab-95-GA-random-3 | HR2_Domain_chainA | −0.9862 | −0.9924 | −0.9930 |
| CoV-Ab-95-GA-random-3 | HR2_Domain_chainB | −0.9856 | −0.9885 | −0.9930 |
| CoV-Ab-95-GA-random-7 | PEPFOLD-model1 | −0.9419 | −0.9425 | −0.9519 |
| CoV-Ab-95-GA-random-7 | HR2_Domain | −2.1792 | −2.1855 | −2.1834 |
| CoV-Ab-95-GA-random-7 | HR2_Domain_chainA | −1.0199 | −1.0191 | −1.0164 |
| CoV-Ab-95-GA-random-7 | HR2_Domain_chainB | −1.0152 | −1.0091 | −1.0228 |
| CoV-Ab-95-seed | PEPFOLD-model1 | −1.0224 | −1.0355 | −1.0262 |
| CoV-Ab-95-seed | HR2_Domain | −2.2681 | −2.2509 | −2.2711 |
| CoV-Ab-95-seed | HR2_Domain_chainA | −1.0745 | −1.0704 | −1.0758 |
| CoV-Ab-95-seed | HR2_Domain_chainB | −1.0715 | −1.0728 | −1.0728 |
| CoV-MUT_AAYL50_21000 | MUT_PEPFOLD-model1 | −0.8903 | −0.8857 | −0.8863 |
| CoV-MUT_AAYL50_21000 | MUT_HR2_Domain_chainB | −1.0202 | −1.0190 | −1.0189 |
| CoV-MUT_Ab-14-SA-PSSM-1 | MUT_PEPFOLD-model1 | −0.7689 | −0.7739 | −0.7708 |
| CoV-MUT_Ab-14-SA-PSSM-1 | MUT_HR2_Domain_chainB | −0.8860 | −0.8866 | −0.8848 |
| CoV-MUT_Ab-14-SA-PSSM-6 | MUT_PEPFOLD-model1 | −0.8073 | −0.8152 | −0.8037 |
| CoV-MUT_Ab-14-SA-PSSM-6 | MUT_HR2_Domain_chainB | −0.9414 | −0.9420 | −0.9360 |
| CoV-MUT_Ab-14-seed | MUT_PEPFOLD-model1 | −0.9085 | −0.8978 | −0.9118 |
| CoV-MUT_Ab-14-seed | MUT_HR2_Domain_chainB | −1.0458 | −1.0486 | −1.0507 |

Supplementary Table 3: Constant-pH Coarse-Grained Modeling for Binding Free Energy Calculations. Estimated binding affinities between antibodies and HR2 peptides, expressed in $k_B$T units. Data were obtained from FORTE simulations conducted at pH 7, 150 mM NaCl, and 298 K. The antigens, named PEPFOLD-model1, HR2 Domain, HR2 Domain chainA, and HR2 Domain chainB, correspond respectively to: (1) the short 50-monomer peptide structure produced by PEP-Fold4, (2) the homotrimer peptide structure provided by PDB ID 8CZI, (3) a single peptide from chain A of the same PDB coordinates, and (4) chain B from the same PDB. The antibody names, extended as -SA-PSSM-1, -SA-PSSM-6, -seed, -GA-random-1, -GA-random-3, and -GA-random-7, are detailed in the text, where further explanations are provided.

| Antibody | Antigen | Estimated Binding Affinity | | | |
|---|---|---|---|---|---|
| | | Replica1 | Replica2 | Replica3 | Average |
| AAYL49_4032 | PEPFOLD-model1 | −1.1203 | −1.1190 | −1.1192 | −1.1195(7) |
| AAYL49_4032 | HR2_Domain | −2.4096 | −2.4107 | −2.4195 | −2.4133(54) |
| AAYL49_4032 | HR2_Domain_chainA | −1.1660 | −1.1633 | −1.1661 | −1.1651(16) |
| AAYL49_4032 | HR2_Domain_chainB | −1.1767 | −1.1600 | −1.1633 | −1.1667(88) |
| AAYL50_21000 | PEPFOLD-model1 | −1.0721 | −1.0704 | −1.0695 | −1.0707(13) |
| AAYL50_21000 | HR2_Domain | −2.3451 | −2.3611 | −2.3522 | −2.3528(80) |
| AAYL50_21000 | HR2_Domain_chainA | −1.1196 | −1.1353 | −1.1278 | −1.1276(79) |
| AAYL50_21000 | HR2_Domain_chainB | −1.1243 | −1.1166 | −1.1366 | −1.1258(101) |
| Ab-14-SA-PSSM-1 | PEPFOLD-model1 | −0.8383 | −0.8412 | −0.8425 | −0.8407(22) |
| Ab-14-SA-PSSM-1 | HR2_Domain | −1.9810 | −1.9570 | −1.9696 | −1.9692(120) |
| Ab-14-SA-PSSM-1 | HR2_Domain_chainA | −0.9072 | −0.9151 | −0.9163 | −0.9129(49) |
| Ab-14-SA-PSSM-1 | HR2_Domain_chainB | −0.9211 | −0.9205 | −0.9121 | −0.9179(50) |
| Ab-14-SA-PSSM-6 | PEPFOLD-model1 | −0.9529 | −0.9584 | −0.9524 | −0.9546(33) |
| Ab-14-SA-PSSM-6 | HR2_Domain | −2.1893 | −2.1905 | −2.1822 | −2.1873(45) |
| Ab-14-SA-PSSM-6 | HR2_Domain_chainA | −1.0236 | −1.0230 | −1.0334 | −1.0267(58) |
| Ab-14-SA-PSSM-6 | HR2_Domain_chainB | −1.0281 | −1.0278 | −1.0271 | −1.0277(5) |
| Ab-14-seed | PEPFOLD-model1 | −1.1157 | −1.1159 | −1.1103 | −1.1140(32) |
| Ab-14-seed | HR2_Domain | −2.4174 | −2.4169 | −2.4096 | −2.4146(44) |
| Ab-14-seed | HR2_Domain_chainA | −1.1506 | −1.1533 | −1.1445 | −1.1495(45) |
| Ab-14-seed | HR2_Domain_chainB | −1.1458 | −1.1472 | −1.1560 | −1.1497(55) |
| Ab-95-GA-random-1 | PEPFOLD-model1 | −0.9011 | −0.8988 | −0.8988 | −0.8996(13) |
| Ab-95-GA-random-1 | HR2_Domain | −2.1154 | −2.1032 | −2.1058 | −2.1081(64) |
| Ab-95-GA-random-1 | HR2_Domain_chainA | −0.9814 | −0.9901 | −0.9810 | −0.9842(51) |
| Ab-95-GA-random-1 | HR2_Domain_chainB | −0.9781 | −0.9856 | −0.9814 | −0.9817(38) |
| Ab-95-GA-random-3 | PEPFOLD-model1 | −0.9090 | −0.9185 | −0.9135 | −0.9137(48) |
| Ab-95-GA-random-3 | HR2_Domain | −2.1195 | −2.1271 | −2.1217 | −2.1228(38) |
| Ab-95-GA-random-3 | HR2_Domain_chainA | −0.9862 | −0.9924 | −0.9930 | −0.9905(38) |
| Ab-95-GA-random-3 | HR2_Domain_chainB | −0.9856 | −0.9885 | −0.9930 | −0.9890(38) |
| Ab-95-GA-random-7 | PEPFOLD-model1 | −0.9419 | −0.9425 | −0.9519 | −0.9454(56) |
| Ab-95-GA-random-7 | HR2_Domain | −2.1792 | −2.1855 | −2.1834 | −2.1827(32) |
| Ab-95-GA-random-7 | HR2_Domain_chainA | −1.0199 | −1.0191 | −1.0164 | −1.0185(18) |
| Ab-95-GA-random-7 | HR2_Domain_chainB | −1.0152 | −1.0091 | −1.0228 | −1.0157(69) |
| Ab-95-seed | PEPFOLD-model1 | −1.0224 | −1.0355 | −1.0262 | −1.0280(67) |
| Ab-95-seed | HR2_Domain | −2.2681 | −2.2509 | −2.2711 | −2.2634(109) |
| Ab-95-seed | HR2_Domain_chainA | −1.0745 | −1.0704 | −1.0758 | −1.0736(28) |
| Ab-95-seed | HR2_Domain_chainB | −1.0715 | −1.0728 | −1.0728 | −1.0724(8) |
| MUT_AAYL50_21000 | MUT_PEPFOLD-model1 | −0.8903 | −0.8857 | −0.8863 | −0.8874(24) |
| MUT_AAYL50_21000 | MUT_HR2_Domain_chainB | −1.0202 | −1.0190 | −1.0189 | −1.0194(7) |
| MUT_Ab-14-SA-PSSM-1 | MUT_PEPFOLD-model1 | −0.7689 | −0.7739 | −0.7708 | −0.7712(25) |
| MUT_Ab-14-SA-PSSM-1 | MUT_HR2_Domain_chainB | −0.8860 | −0.8866 | −0.8848 | −0.8858(9) |
| MUT_Ab-14-SA-PSSM-6 | MUT_PEPFOLD-model1 | −0.8073 | −0.8152 | −0.8037 | −0.8087(59) |
| MUT_Ab-14-SA-PSSM-6 | MUT_HR2_Domain_chainB | −0.9414 | −0.9420 | −0.9360 | −0.9398(32) |
| MUT_Ab-14-seed | MUT_PEPFOLD-model1 | −0.9085 | −0.8978 | −0.9118 | −0.9060(73) |
| MUT_Ab-14-seed | MUT_HR2_Domain_chainB | −1.0458 | −1.0486 | −1.0507 | −1.0484(25) |

Supplementary Table 5: Statistical significance of biophysical properties for antibodies from different generation approaches comparing with the training antibodies. P-values> 0.05 are highlighted in red.

| **Method** | **Instability** | | **Isoelectric_point** | | **mean_Hydrophobicity** | | **Helix_fraction** | | **Turn_fraction** | | **Sheet_fraction** | |
|---|---|---|---|---|---|---|---|---|---|---|---|---|
| | t-statistics | p-value | t-statistics | p-value | t-statistics | p-value | t-statistics | p-value | t-statistics | p-value | t-statistics | p-value |
| Ab-14-SA-random | 8.618 | 3.53E-10 | 22.732 | 6.35E-23 | 2.838 | 8.09E-03 | 51.424 | 4.25E-34 | -38.108 | 3.60E-24 | -166.062 | 1.38E-55 |
| Ab-14-SA-pssm | 5.917 | 1.64E-06 | 30.655 | 2.58E-28 | 10.637 | 5.75E-12 | 36.048 | 1.00E-29 | -32.199 | 1.41E-24 | -131.73 | 5.28E-48 |
| Ab-14-SA-abmodel | 6.129 | 1.29E-06 | 20.536 | 1.40E-20 | 1.422 | 1.64E-01 | 31.904 | 2.93E-25 | -26.92 | 1.09E-25 | -147.999 | 3.83E-53 |
| Ab-14-GA-random | 24.708 | 5.69E-17 | 3.507 | 1.40E-03 | -3.083 | 5.04E-03 | 44.323 | 3.44E-34 | -33.173 | 6.84E-27 | -186.376 | 2.95E-54 |
| Ab-14-GA-pssm | 18.347 | 9.55E-18 | 12.046 | 1.36E-12 | -3.008 | 6.27E-03 | 60.958 | 1.92E-36 | -36.774 | 2.14E-25 | -205.893 | 5.39E-49 |
| Ab-14-GA-abmodel | 3.752 | 6.00E-04 | 6.595 | 2.59E-07 | -3.528 | 1.80E-03 | 43.21 | 2.31E-33 | -34.08 | 9.12E-28 | -177.822 | 5.88E-56 |
| Ab-91-SA-random | 3.909 | 4.00E-04 | -3.161 | 4.38E-03 | 16.681 | 1.05E-17 | 60.039 | 3.74E-34 | -34.927 | 2.81E-30 | -105.486 | 6.96E-41 |
| Ab-91-SA-pssm | 18.274 | 2.12E-20 | -4.442 | 2.55E-04 | -2.284 | 3.05E-02 | 39.041 | 3.31E-32 | -18.976 | 5.75E-21 | -134.134 | 1.39E-49 |
| Ab-91-SA-abmodel | 15.439 | 1.87E-15 | -5.263 | 3.83E-05 | -6.888 | 1.53E-07 | 28.637 | 4.59E-27 | -20.019 | 1.27E-20 | -219.462 | 5.95E-40 |
| Ab-91-GA-random | 15.556 | 4.58E-18 | -3.855 | 9.90E-04 | 8.583 | 5.09E-09 | 44.861 | 5.11E-34 | -38.288 | 2.27E-27 | -144.071 | 3.65E-53 |
| Ab-91-GA-pssm | 3.302 | 2.39E-03 | -3.464 | 2.10E-03 | 6.422 | 2.01E-06 | 48.2 | 2.60E-34 | -34.058 | 3.86E-29 | -168.965 | 2.33E-51 |
| Ab-91-GA-abmodel | 6.138 | 5.35E-07 | -5.737 | 1.51E-05 | 4.506 | 6.85E-05 | 21.034 | 7.07E-20 | -14.972 | 3.19E-17 | -156.636 | 1.16E-53 |
| Ab-95-SA-random | 3.854 | 6.36E-04 | 2.028 | 5.51E-02 | -13.14 | 7.27E-15 | 39.733 | 3.48E-28 | -42.995 | 4.72E-29 | -166.255 | 2.10E-51 |
| Ab-95-SA-pssm | 10.847 | 8.51E-13 | -1.059 | 2.97E-01 | -3.587 | 9.40E-04 | 31.034 | 5.73E-28 | -46.273 | 7.81E-35 | -132.536 | 1.61E-51 |
| Ab-95-SA-abmodel | 3.784 | 7.17E-04 | 8.000 | 1.49E-07 | -20.414 | 6.07E-22 | 16.3 | 1.92E-18 | -39.161 | 1.07E-27 | -142.658 | 1.95E-53 |
| Ab-95-GA-random | -3.756 | 8.34E-04 | 1.000 | 3.24E-01 | 2.299 | 2.88E-02 | 23.168 | 7.43E-22 | -47.178 | 7.68E-33 | -172.32 | 2.13E-35 |
| Ab-95-GA-pssm | 1.246 | 2.23E-01 | 0.244 | 8.09E-01 | -2.782 | 8.82E-03 | 36.903 | 9.72E-31 | -40.03 | 1.27E-30 | -146.395 | 8.32E-54 |
| Ab-95-GA-abmodel | 7.278 | 5.38E-08 | 4.465 | 2.28E-04 | -1.194 | 2.40E-01 | 31.113 | 1.66E-28 | -45.876 | 5.89E-29 | -155.191 | 1.32E-51 |
